# Structural and Biological Evaluations of a Non-Nucleoside STING Agonist Specific for Human STING^A230^ Variants

**DOI:** 10.1101/2023.07.02.547363

**Authors:** Zhichao Tang, Junxing Zhao, Ying Li, Shallu Tomer, Manikandan Selvaraju, Nicholas Tien, Diyun Sun, David K. Johnson, Anjie Zhen, Pingwei Li, Jingxin Wang

## Abstract

Previously we identified a non-nucleotide tricyclic agonist BDW568 that activates human STING (stimulator of interferon genes) gene variant containing A230 in a human monocyte cell line (THP-1). STING^A230^ alleles, including HAQ and AQ, are less common STING variants in human population. To further characterize the mechanism of BDW568, we obtained the crystal structure of the C-terminal domain of STING^A230^ complexed with BDW-OH (active metabolite of BDW568) at 1.95 Å resolution and found the planar tricyclic structure in BDW-OH dimerizes in the STING binding pocket and mimics the two nucleobases of the endogenous STING ligand 2’,3’-cGAMP. This binding mode also resembles a known synthetic ligand of human STING, MSA-2, but not another tricyclic mouse STING agonist DMXAA. Structure-activity-relationship (SAR) studies revealed that all three heterocycles in BDW568 and the S-acetate side chain are critical for retaining the compound’s activity. BDW568 could robustly activate the STING pathway in human primary peripheral blood mononuclear cells (PBMCs) with STING^A230^ genotype from healthy individuals. We also observed BDW568 could robustly activate type I interferon signaling in purified human primary macrophages that were transduced with lentivirus expressing STING^A230^, suggesting its potential use to selectively activate genetically engineered macrophages in macrophage-based approaches, such as chimeric antigen receptor (CAR)-macrophage immunotherapies.

## Introduction

Cyclic GMP-AMP synthase (cGAS)-STING pathway is crucial for recognizing self or foreign double-stranded DNA and activating type I interferon (IFN-I) signaling via the interferon regulatory factor 3 (IRF3) axis.^1,2^ This pathway is indispensable for the innate immune responses in mammalian cells in events of bacteria or DNA virus infections.^3,4^ Pharmacological STING activation has been demonstrated as a potential promising approach for the treatment of cancer and virus infection,^5–10^ as well as vaccine adjuvants.^11–13^ Human STING has genetic polymorphisms. Recent genetic analysis demonstrated that the most common STING allele is R71-G230-R293 (RGR) in 59.2% of human population, followed by H71-A230-Q293 (HAQ) which occurs in 20.4%, H232 in 13.7%, A230-Q293 (AQ) in 5.2%, and Q293 in 1.5%.^14^ The STING-HAQ variant was found negatively affect innate immunity in humans compared to the most common STING allele (RGR),^14–17^ while others claimed that the difference is insignificant.^18^

To our knowledge, almost no reported human STING agonists can discriminate against STING variants except for BDW568 (Figure 1).^5–7,19,20^ BDW568 is a newly identified STING agonist by our laboratory which can selectively activate STING^A230^ variants,^21^ including HAQ and AQ in human population. The activity of BDW568 against G230 STING was not observed, suggesting a nearly complete selectivity in naturally occurring STING variants.^21^ BDW568 is a methyl ester and can be quickly hydrolyzed after cellular uptake by cytosolic carboxylesterase 1 (CES1) to yield the active metabolite BDW-OH (Figure 1).^21^ Here, we reported the crystal structure of the STING^A230^ C-terminal domain (CTD)–BDW-OH complex and structural-activity-relationship studies. We also validated the activity of BDW568 in primary human cells from healthy individuals with endogenous STING^A230^ allele or transduced exogenous STING^A230^ in purified primary macrophages.

**Figure 1.**
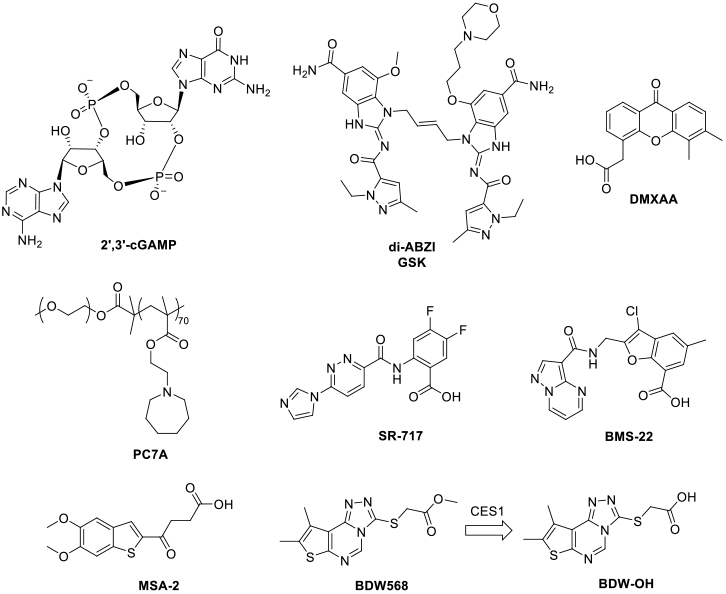
Structures of natural and synthetic STING agonists.

## Results and Discussion

### Crystal structure of STING^A230^ CTD – BDW-OH complex

To acquire the structural information of STING^A230^–BDW-OH complex, we expressed small ubiquitin-related modifier (SUMO) fusion of human STING^A230^ CTD (residue 155–341) and co-crystalized the protein with purified BDW-OH or with STING endogenous ligand, 2’,3’-cyclic GMP-AMP (cGAMP). The crystal structures of the STING^A230^ CTD complexed with BDW-OH and 2’,3’-cGAMP were obtained at 1.95 and 2.01 Å resolution, respectively (Table S1, PDB: 8T5K, 8T5L). Comparing these two structures, we found the overall conformation of STING^A230^–BDW-OH complex is similar to that of STING^A230^–2’,3’-cGAMP complex (Figure 2). The STING^A230^ CTD forms a butterfly-shaped dimer. Unlike the apo form of STING, in which the two “wings” from the two STING monomers are wide open, the STING–ligand complexes adopt a closed conformation (Figures 2A & 2B).

**Figure 2.**
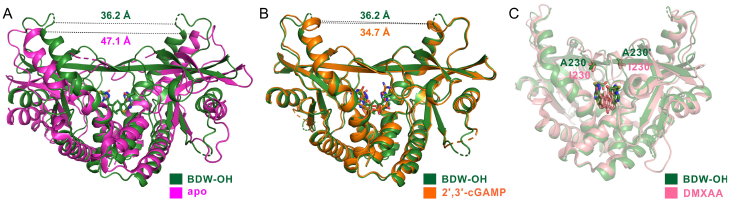
Overlay of STING^A230^–BDW-OH complex (PBD: 8T5K) with (A) STING apo form (PBD: 4EMU),^22^ (B) STING^A230^– 2’,3’-cGAMP complex (PBD: 8T5L), and STING^I230^–DMXAA complex (PDB: 4QXP).^23^ The distance between residues H185 from both monomers is shown to illustrate open (∼47 Å) and closed (34–37Å) conformations.

As expected, two molecules of BDW-OH occupy the 2’3’-cGAMP binding pocket in STING^A230^ homodimer in a symmetric manner. The carboxylic acid group of BDW-OH engages R238/R232 of STING via hydrogen bonding and the pyrimidine ring of BDW-OH also forms a hydrogen bond with T263 (Figure 3A). In addition, the triazole ring of BDW-OH forms a π-π stacking with Y167 of STING (Figure 3A). These interactions between the ligand and STING were also observed in structures of STING complexed with 2’,3’-cGAMP (Figure 3B) and MSA-2 (PDB: 6UKM),^5^ respectively. Overlay of BDW-OH and 2’,3’-cGAMP revealed that the tricyclic structures of BDW-OH are in the same plane as the two nucleobases of 2’,3’-cGAMP, maintaining the π-π stacking with Y167 (Figure 3C). Similarly, the tricyclic structure of BDW-OH also overlaps well with the benzothiophene ring of MSA-2. Importantly, both BDW-OH and MSA-2 use the carboxylic acid to tackle R238/R232 in similar conformations (Figure 3D). Therefore, we concluded that that binding mode of STING^A230^ and BDW-OH resembles the STING–MSA-2 complex. We previously speculated that BDW-OH binds to STING^A230^ in a similar binding mode as that of a mouse STING-specific tricyclic ligand, DMXAA, and an artificial STING mutant with I230. Interestingly, although the DMXAA binds in the same pocket as BDW-OH (Figure 2C), its tricyclic ring does not in the same plane as that of BDW-OH (Figure 3E). The orientation of the carboxylic acid group of DMXAA is also markedly different from that of BDW-OH in the bound conformation for the interaction with R238 of STING (Figure 3E).

**Figure 3.**
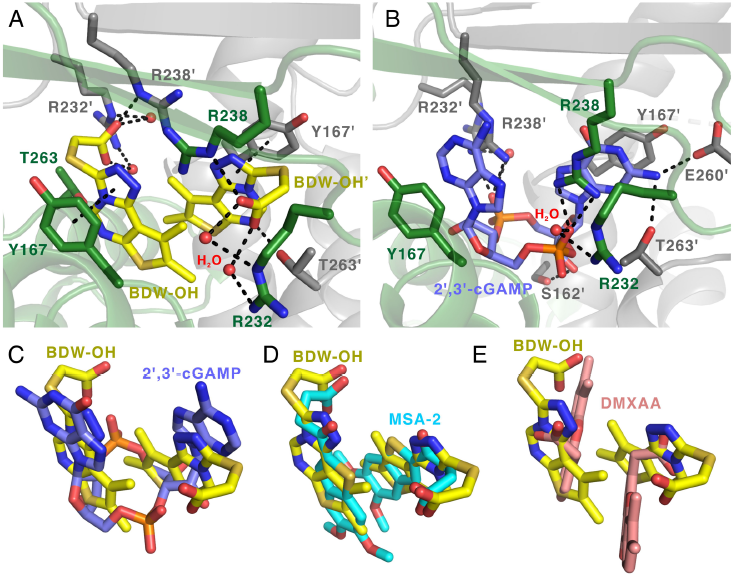
Interactions between STING^A230^ CTD and (A) BDW-OH (PDB: 8T5K) and (B) 2’,3’-cGAMP (PDB: 8T5L). Superimposition of the bound structures of BDW-OH and (C) 2’,3’-cGAMP, (D) MSA-2 (PDB: 6UKM),^5^ and (E) DMXAA (PDB: 4QXP).^23^

Residue 230 of human STING is located in the lid region of a STING dimer (Figure 2C).^23^ We confirmed from the crystal structure that A230 does not interact with BDW-OH. The lid region is a four-stranded, antiparallel β-sheet that covers the binding pocket of STING. It was demonstrated that single-point mutation in the lid region can control the pharmacological activation of STING. For example, G230I mutation in human STING sensitized the mouse STING-specific DMXAA to bind.^23^ Similarly, I229G or I229A mutation in mouse STING desensitized the DMXAA binding (residue 229 in mouse STING is homologous to residue 230 in humans).^23^ In human STING, because residue 230 does not interact with the BDW-OH directly, we further hypothesized that the residue 230 gates the exit tunnel of the ligand as a mechanism of ligand recognition. As such, bulkier amino acid residues (i.e., alanine or isoleucine) can probably retain ligands better than glycine residue.^21^

### Chemical modifications on the tricyclic structure

To further probe the interactions between BDW-OH and STING^A230^, we performed structure-activity-relationship (SAR) studies and used an interferon-sensitive response element (ISRE) reporter gene assay in THP-1 cells to evaluate the compounds’ activity.^21^ The ISRE response to IFN-I signaling is a key downstream event of STING activation and is routinely used to quantify potency for STING agonists.^6^ The genotype of STING in THP-1 cells is homozygous STING^HAQ^.^14^ We previously demonstrated that STING activation in THP-1 cells using BDW568 is solely dependent on A230, but not H71 or Q293.^21^ The crystal structure of STING^A230^-CTD–BDW-OH showed that there is a small unoccupied cavity on the dimethyl thiophene side in BDW568 (Figure 3A). To probe the size of the cavity, we linked the 4,5-dimethyl groups on the thiophene ring (A) with one (**1**) or two carbons (**2**) (Table 1). The reactivity (illustrated as half maximal effective concentrations, or EC_50_’s) of compounds **1** was retained while compound **2** was inactive (Table 1), implying the cavity in the binding pocket can only accommodate one carbon. An isomer of BDW568, having an ethyl group at C5 and hydrogen at C4 (**3**) is also active. In contrast, removal of the dimethyl groups (**4**) significantly weakened the activity (Table 1). On the other hand, substitution of the methyl group at C4 position of the thiophene with a bulkier group, such as phenyl group (**5**), completely abrogated the compound’s activity, confirming that the space close to the thiophene ring (A) is limited. Similarly, substitution of either of the two methyl groups with a methoxy group (**6, 7**), or replacing the 5-methyl group on the thiophene ring into a bromo group (**8**), all significantly reduced the activity (Table 1).

**Table 1.** ISRE activation activities of BDW568 analogs in THP-1 cells.

| Compound | A/B/C rings | EC<sub>50</sub> (μM) |
| --- | --- | --- |
| BDW568 |  | 5.7 ± 1.3 |
| 1 |  | 7.3 ± 0.9 |
| 2 |  | NA |
| 3 |  | 6.9 ± 2.0 |
| 4 |  | NA |
| 5 |  | NA |
| 6 |  | 23.2 ± 16.2 |
| 7 |  | NA |
| 8 |  | 11.4 ± 6.4 |
| 9 |  | NA |
| 10 |  | NA |
| 11 |  | NA |
| 12<sup>21</sup> |  | NA |
| 13<sup>21</sup> |  | NA |
| 14<sup>21</sup> |  | NA |
| 15 |  | NA |

NA = not active

In addition, we found the skeleton of the tricyclic structure cannot be altered for STING activation. For example, replacing the thiophene ring (A) with pyrrole (**9**), furan (**10**) or dimethoxybenzene (**11**) completely abrogates the compounds’ activity (Table 1). This is consistent with our previous report, in which we demonstrated that replacing the pyrimidine ring (B) with pyridine (**12**) or 2-methyl pyrimidine (**13**), or triazole ring (C) with imidazole (**14**) all failed to retain the activity.^21^ Similarly, breaking the triazole ring (C) by using an N-carbamoylacetamide group (**15**) was also intolerable (Table 1).

### Chemical modifications on the S-acetate side chain

BDW568 is required to be hydrolyzed by CES1 to yield the carboxylic acid metabolite to interact with STING^A230^.^21^ We tested other esters, such as ethyl (**16**) and isopropyl esters (**17**) and observed a slightly weakened activity (Table 2). Further increasing the bulkiness of the ester to tert-butyl ester (**18**) or changing the linear ester into a lactone (**19**) completely diminished the activity probably because these esters can no longer be recognized by CES1. We also synthesized several cell permeable carboxylic isosteres, such as tetrazole (**20**) and anisole (**21**)^24^ and observed that none of these isosteres could retain the activity. We concluded that a carboxylic acid group in BDW568 is required for STING binding. The side chain also needs to maintain a proper length to engage the R238 residue in STING, since addition (**22**) or reduction (**23**) of the chain length completely diminished the activity (Table 2). Interestingly, the thioether group in the side chain is also crucial for STING activation. We replaced the S into O (**24**) and CH_2_ (**25**), or oxidized the S into sulfoxide (**26**), and found all these the alternations inactivated the compounds (Table 2).

**Table 2.** ISRE activation activities of BDW568 analogs in THP-1 cells.

| Compound | Side chain | EC<sub>50</sub> (μM) |
| --- | --- | --- |
| BDW568 |  | 5.7 ± 1.3 |
| 16 |  | 8.8 ± 0.9 |
| 17<sup>21</sup> |  | 13.4 ± 0.5 |
| 18<sup>21</sup> |  | NA |
| 19 |  | NA |
| 20<sup>21</sup> |  | NA |
| 21 |  | NA |
| 22 |  | NA |
| 23 |  | NA |
| 24 |  | NA |
| 25 |  | NA |
| 26 |  | NA |

NA = not active

### Compound synthesis

Compounds with modifications on tricyclic rings (**1**–**11** and **15**) were synthesized as outlined in Scheme 1, which is similar to the synthetic route for BDW568.^21^ Starting with substituted pyrimidinone (S1), the chloro-substituted intermediate S2 was obtained by refluxing in POCl_3_ with high yield. Subsequent S_N_2 reaction in hydrazine hydrate yielded the key intermediate S3 which also served as a starting material for compounds with O/CH_2_ side chain modifications (see Scheme 2). S3 then underwent ring-closing rection with CS_2_/KOH to form the triazole ring and afforded S4. The reaction between S4 and methyl bromoacetate in DMF/DIPEA occurred rapidly and usually completed within several minutes. Compound **15** was obtained by a single urea-formation reaction using triphosgene as the coupling reagent.

**Scheme 1.**
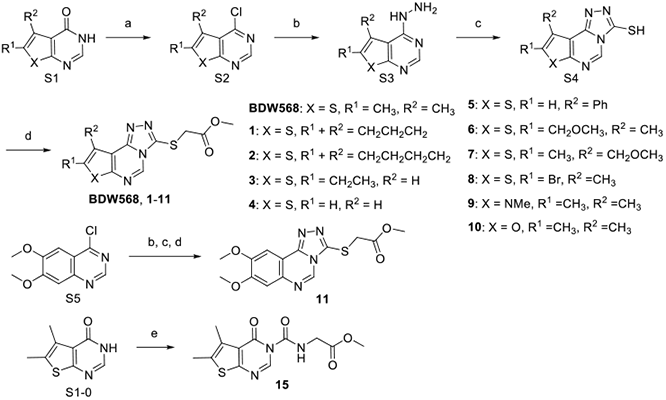
Synthesis of compounds with modifications on the tricyclic rings.^*a*^ ^*a*^ Reagents and conditions: (a) POCl_3_, reflux, 1 h; (b) 65% hydrazine, 100 °C, 30 min; (c) CS_2_, KOH, EtOH, 90 °C, 1 h; (d) methyl bromoacetate, DIPEA, DMF, r.t., 30 min; (e) methyl 2-aminoacetate, triphosgene, CH_2_Cl_2_, r.t., overnight (DMF = dimethylformamide DIPEA = *N,N*-diisopropylethylamine, r.t. = room temperature).

**Scheme 2.**
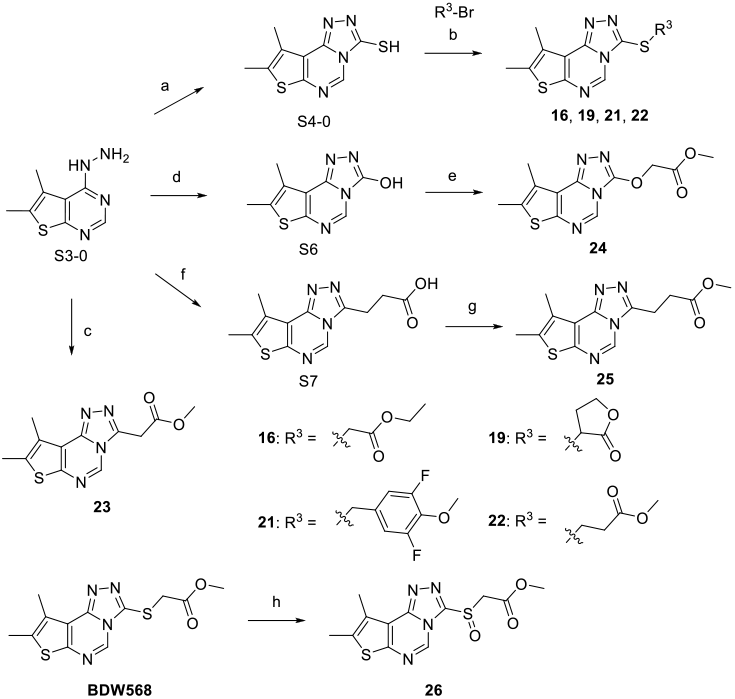
Synthesis of compounds with modifications on the side chain.^*a*^ ^*a*^Reagents and conditions: (a) CS_2_, KOH, EtOH, 90 °C, 1 h; (b) R^3^-Br, DIPEA, DMF, r.t., 30 min; (c) dimethyl malonate, microwave, 180 °C, 45 min; (d) 1, 1’-carbonyldiimidazole, MeCN, 90 °C, 1 h; (e) NaH, DMF, 0 °C–r.t., 30 min; (f) succinic anhydride, DMF, 80 °C, overnight; (g) MeOH, concentrated H_2_SO_4_, reflux, 2 h; (h) mCPBA, CH_2_Cl_2_, r.t., 2 h.

Compounds with side chain modifications can be prepared as shown in Scheme 2. Intermediate S4 and substituted alky bromides were used to synthesize compounds **16, 19, 21** and **22** with a S-linker. Intermediate S3 was used in different conditions to form the desired ring closed product. A single reaction of S3 in dimethyl malonate under microwave irradiation and high temperature directly yielded target compound **23**. 1,1’-carbonyldiimidazole (CDI) was used in the ring-closing reaction of intermediate S6, and subsequent S_N_2 reaction in NaH/DMF gave desired product **24**. Reaction between succinic anhydride and S3 could produce the ring closed intermediate S7 in carboxylic acid form which was then refluxed in methanol with concentrated H_2_SO_4_ as catalyst to afford compound **25**. The sulfur oxidized compound **26** was obtained from BDW568 by using mCPBA as oxidant.

### Biological activities in human primary cells and CAR-macrophages

Next, we set out to explore the potential application of BDW568 in immunotherapy. In the first step, we confirmed the activity of BDW568 in THP-1 cells by real-time quantitative PCR (RT-qPCR) and found that THP1 cells have elevated expression of interferon stimulated genes (ISGs), including interferon-induced GTP-binding protein (MX1) and 2’,5’-oligoadenylate synthetase 1 (OAS1) after BDW568 stimulation (Figure 4A). We selected the MX1 gene as a marker to quantify IFN-I activation in following experiments. Then, to examine the activity of BDW568 in primary human cells, we collected PBMCs from 15 healthy donors and measured MX1 expression in the presence of BDW568 and 2’,3’-cGAMP. The PBMCs from donor 9 responded to BDW568, while all donors responded to 2’,3’-cGAMP (Figure 4B). We genotyped donor 9 and confirmed that it carries homozygous STING^A230^. The rest 14 donors do not have a STING^A230^ allele.

**Figure 4.**
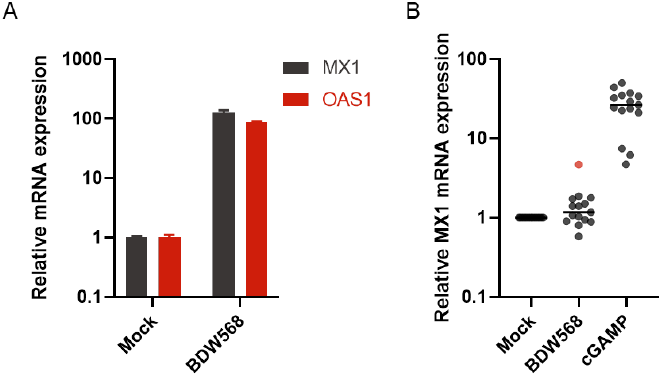
BDW568 stimulates ISG expression in THP-1 cells and primary PBMCs that carry STING^230A^. (A) THP-1 cells were stimulated with either vehicle or BDW568 (50 μM) for 6 h. Afterwards, cells were harvested, and RT-qPCR were performed to measure expression level of MX1 and OAS1. (B) Primary PBMCs were collected from 15 independent healthy donors, cells were then stimulated with BDW568 (50 μM) and 2’,3’-cGAMP (1 μg/ml) for 6 h. Afterwards, the cells were harvested, and RT-qPCR were performed to measure MX1 expression level. Donor 9 (red) showed significant elevation of MX1 after BDW568 stimulation.

Encouraged by the promising results in human primary cells, we envision that BDW568 can be a useful probe to selectively activate the STING pathway in genetically engineered immunogenic cells that carry the STING^A230^ allele. To examine if we can genetically engineer cells with STING^A230^ to respond to BDW568, we generated lentivirus that will allow overexpression of full-length STING^A230^ and a marker protein, which is a truncated inactive EGF receptor (EGRt) (Figure 5). Monocyte derived macrophages were generated from 3 dependent healthy donors with STING^G230^ alleles and transduced with EGFRt-STING^A230^ lentivirus. 3 days after transduction, cells were stimulated with vehicle or BDW568. As expected, no elevation of the MX1 gene was observed in mock transduced cells or untransduced cells after BDW568 stimulation. In contrast, macrophages expressing EGFRt-STING^A230^ showed a 10-fold increase in MX1 transcription (Figure 5), implying a robust selectivity for pharmacologically activation of STING^A230^ engineered macrophages without affecting the STING^G230^ cells. As such, STING^A230^-specific agonist can be potentially used in STING^G230^ patients to selectively activate engineered cells in cellular therapy, such as chimeric antigen receptor (CAR)-macrophages, which recently showed promising activity to treat solid tumors.^25^

**Figure 5.**
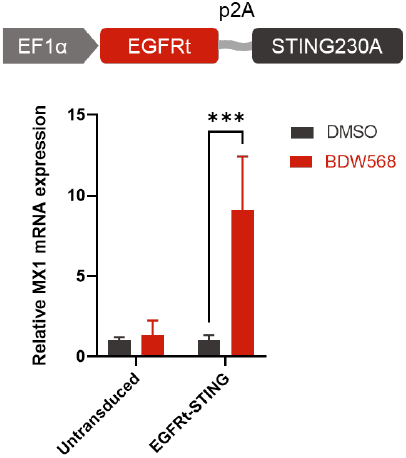
BDW568 stimulation showed upregulated expression of ISG MX1 in primary macrophages transduced with lentivirus STING^230A^ as compared to untransduced macrophages. CD14^+^ monocytes were differentiated into macrophages in the presence of M-CSF. Cells were transduced with lentivirus FG12-EF1p-EGFRt-p2A-STING^230A^ or mock transduced and stimulated with DMSO (vehicle) or BDW568 for 6 h. ISG MX1 gene expression level was measured by RT-PCR (n = 3, ***P < 0.0005).

## Conclusion

In summary, we obtained the crystal structure of STING^A230^ with a newly reported STING agonist, BDW568. In STING^A230^–BDW-OH (active metabolite of BDW568) complex The STING dimer adopts a “closed” conformation, which is almost identical to that of STING^A230^–2’,3’-cGAMP (endogenous ligand) complex. By comparing the structures of STING^A230^–BDW-OH complex with other reported structures, we concluded that the binding mode of BDW-OH is similar to a known synthetic STING ligand, MSA-2. SAR studies demonstrated that the skeleton of all three heterocycles is crucial for BDW568’s activity and cannot be extensively modified. We also found that the S-acetate side chain is essential to maintain the activity. We confirmed that the BDW568’s selectivity in cells with STING^A230^ allele was observed in primary PBMCs. Importantly, BDW568 can robustly and selectively activate macrophages that were transduced with STING^A230^. Therefore, this compound may be used to selectively activate STING^A230^ engineered macrophages for macrophage-based immunotherapies, including CAR-macrophage therapies.

## Experimental Section

### Crystallography

#### Protein expression and purification

The gene encoding human STING (155–341) 230A/232R was cloned into a home-modified pET28a vector with a *N*-terminal His_6_ and SUMO tag. The protein was expressed in *Escherichia coli* BL21 (DE3) cells induced with 0.4 mM isopropyl β-d-1-thiogalactopyranoside (IPTG) overnight at 16°C. The protein was purified by Ni-NTA agarose. The N terminal tag was cleaved by SUMO protease and removed by a Ni-NTA column. And protein was further purified by a HiLoad 16/600 Superdex 200 column in a buffer containing 20 mM Tris-HCI pH 7.5, 150 mM NaCl and concentrated to 13 mg/ml for crystallization.

#### Crystallization and structure determination

Purified STING (155–341) 230A/232R was mixed with BDW-OH at the molar ratio of 1:5. Crystallization screening was performed by hanging drop vapor diffusion method at 4°C using Index and PEGRx kits from Hampton Research. Crystals were grown in a reservoir solution containing 0.2 M ammonium sulfate, 0.1 M HEPES pH 7.5, 25% polyethylene glycol 3350. After two days, crystals were harvested, cryoprotected in reservoir solution containing 30% sucrose, and flash frozen in liquid nitrogen. Crystallization of STING 230A/232R in complex with 2’,3’-cGAMP was performed at the same condition. Diffraction data were collected using a Rigaku MicroMax-007 HF generator with a RAXIS IV^2+^ detector at home. The data were processed with iMosflm^26^ in the CCP4 package. The structures were determined by molecular replacement using the STING CTD structure (PDB ID: 4KSY)^27^ as a searching model in Phenix.^28^ The structures were rebuilt with coot^29^ and refined by Phenix. Coordinates for the two crystal structures have been deposited into the RCSB Protein Data Bank (PDB) with STING–BDW-OH under accession number 8T5K, and STING– 2’,3’-cGAMP complex under accession number 8T5L.

#### Chemistry

Reagents and solvents were purchased from commercial sources (Fisher, Sigma-Aldrich, Combi-Blocks and Suzhou Medinoah Ltd) and used as received. Reactions were tracked by TLC (Silica gel 60 F_254_, Merck) and Waters ACQUITY UPLC-MS system (ACQUITY UPLC H Class Plus in tandem with QDa Mass Detector). Intermediates and products were purified by a Teledyne ISCO Combi-Flash system using prepacked SiO_2_ cartridges. NMR spectra were acquired on a Bruker AV400 or AV500 instrument (500 MHz for ^1^H NMR, 126 MHz for ^13^C NMR). ^13^C shifts were obtained with ^1^H decoupling. MestReNova 14.0.1 developed by MESTRELAB RESEARCH was used for NMR data processing. MS-ESI spectra were recorded on Waters Qda Mass Detector. The UPLC-MS was performed on a Waters BEH C18 column (2.1 mm × 50 mm, 1.7 μm) with peak detection at UV 254 nm (mobile phase: acetonitrile and 0.1% formic acid in water; gradient: 0–5 min, 2–98% acetonitrile). Purities of final compounds were assessed by UPLC-MS. All compounds are > 95% pure by UPLC analysis.

#### General Procedures

1. Compound S1 (0.5 g, 1 equiv.) in POCl_3_ (4 mL) was refluxed at 120 °C for 1 h or until the reaction completed. POCl_3_ was evaporated in vacuo and the residue was dissolved in ethyl acetate. After washing with saturated NaHCO_3_ solution, the organic phase was collected and dried with Na_2_SO_4_ and concentrated in vacuo to furnish compound S2. The crude product was purified by silica-gel column chromatography using 0–10% ethyl acetate in hexanes.
2. Compound S2 (0.4 g, 1 equiv.) in 65% hydrazine hydrate (4 mL) was heated at 100 °C for 30 min. The solid formed was filtered and washed with water to remove excess hydrazine. The water residue was removed by co-evaporating with toluene to afford compound S3 as yellow solid which was used in the next step without further purification.
3. A solution of KOH (1.3 equiv.) and CS_2_ (4 equiv.) in ethanol was added dropwise to the solution of compound S3 (0.1 g, 1equiv) in ethanol. The reaction mixture was then heated to 90 °C for 1 h. The solvent was removed in vacuo and the residue was dissolved in water and acidified with 1N HCl. The precipitate was filtered, washed with water, and dried. The crude solid was heated and re-crystallized in ethanol to give compound S4.
4. Compound S4 (50 mg, 1 equiv.) in DMF was added the respective alkyl bromide (1.5 equiv.) followed by N, N-diisopropylethylamine (DIPEA, 3 equiv.) and the reaction mixture was stirred at room temperature for 30 min. After completion, the reaction mixture was added water and extracted with ethyl acetate. The organic layer was washed with water followed by brine and was dried with Na_2_SO_4_. Ethyl acetate was removed under vacuum and the crude product was purified by silica-gel column chromatography with 0– 100 % ethyl acetate in hexanes to afford the final product.

Compounds **1** to **11** were synthesized following General Procedures 1 to 4 by using respective starting material S1.

#### Methyl 2-((9,10-dihydro-8*H*-cyclopenta[4,5]thieno[3,2-*e*][1,2,4]triazolo[4,3-*c*]pyrimidin-3-yl)thio)acetate (1)

Light yellow solid; yield: 45%; purity: >95%. ^1^H NMR (500 MHz, DMSO_d6_) δ 9.16 (s, 1H), 4.13 (s, 2H), 3.61 (s, 3H), 3.14 – 3.10 (m, 4H), 2.58 – 2.51 (m, 2H); ^13^C NMR (126 MHz, DMSO_d6_) δ 169.5, 155.5, 147.3, 144.6, 140.8, 138.3, 134.4, 114.8, 53.0, 36.2, 30.1, 28.8, 28.1. Mass ESI [M+H]^+^ = 320.93.

#### Methyl 2-((8,9,10,11-tetrahydrobenzo[4,5]thieno[3,2-*e*][1,2,4]triazolo[4,3-*c*]pyrimidin-3-yl)thio)acetate (2)

Light yellow solid; yield: 41%; purity: >95%. ^1^H NMR (500 MHz, DMSO_d6_) δ 9.50 (s, 1H), 4.22 (s, 2H), 3.70 (s, 3H), 3.00 (t, *J* = 10 Hz, 2H), 2.91 (t, *J* = 10 Hz, 2H), 1.91 – 1.88 (m, 4H); ^13^C NMR (126 MHz, DMSO_d6_) δ 169.5, 165.4, 153.7, 149.2, 138.8, 136.7, 128.8, 119.1, 53.0, 33.4, 25.3, 25.2, 22.9, 22.1. Mass ESI [M+H]^+^= 334.96.

#### Methyl 2-((8-ethylthieno[3,2-*e*][1,2,4]triazolo[4,3-*c*]pyrimidin-3-yl)thio)acetate (3)

Off-white solid; yield: 50%; purity: >95%. ^1^H NMR (500 MHz, DMSO_d6_) δ 9.26 (s, 1H), 7.69 (t, *J* = 1.2 Hz, 1H), 4.19 (s, 2H), 3.67 (s, 3H), 3.09 (qd, *J* = 7.5, 1.2 Hz, 2H), 1.42 (t, *J* = 7.5 Hz, 3H); ^13^C NMR (126 MHz, DMSO_d6_) δ 169.5, 151.1, 150.8, 147.2, 141.1, 135.1, 118.7, 116.5, 53.0, 36.2, 24.0, 15.9. Mass ESI [M+H]^+^ = 308.94.

#### Methyl 2-(thieno[3,2-*e*][1,2,4]triazolo[4,3-*c*]pyrimidin-3-ylthio)acetate (4)

Light yellow solid; yield: 54%; purity: >95%. ^1^H NMR (500 MHz, DMSO_d6_) δ 9.27 (s, 1H), 8.10 (d, *J* = 5.8 Hz, 1H), 7.88 (d, *J* = 5.7 Hz, 1H), 4.15 (s, 3H), 3.62 (s, 3H); ^13^C NMR (126 MHz, DMSO_d6_) δ 169.5, 152.9, 147.5, 141.4, 135.9, 129.9, 120.3, 118.6, 53.0, 36.2. Mass ESI [M+H]^+^ = 280.93.

#### Methyl 2-((9-phenylthieno[3,2-*e*][1,2,4]triazolo[4,3-*c*]pyrimidin-3-yl)thio)acetate (5)

Light yellow solid; yield: 55%; purity: >95%. ^1^H NMR (500 MHz, DMSO_d6_) δ 9.32 (s, 1H), 8.09 (s, 1H), 7.88 – 7.86 (m, 2H), 7.53 – 7.50 (m, 2H), 7.48 – 7.44 (m, 1H), 4.14 (s, 2H), 3.62 (s, 3H); ^13^C NMR (126 MHz, DMSO_d6_) δ 169.4, 153.6, 147.8, 141.4, 137.0, 136.1, 134.7, 129.6, 128.6, 126.0, 116.6, 53.0, 36.1. Mass ESI [M+H]^+^ = 356.97.

#### Methyl 2-((8-(methoxymethyl)-9-methylthieno[3,2-*e*][1,2,4]triazolo[4,3-*c*]pyrimidin-3-yl)thio)acetate (6)

Off-white solid; yield: 52%; purity: >95%. ^1^H NMR (500 MHz, DMSO_d6_) δ 9.22 (s, 1H), 4.76 (s, 2H), 4.14 (s, 2H), 3.62 (s, 3H), 3.39 (s, 4H), 2.69 (s, 3H); ^13^C NMR (126 MHz, DMSO_d6_) δ 169.0, 150.3, 147.4, 140.5, 137.4, 135.1, 128.3, 118.8, 66.4, 57.9, 52.5, 35.7, 13.3. Mass ESI [M+H]^+^ = 338.95.

#### Methyl 2-((9-(methoxymethyl)-8-methylthieno[3,2-*e*][1,2,4]triazolo[4,3-*c*]pyrimidin-3-yl)thio)acetate (7)

Off-white solid; yield: 62%; purity: >95%. ^1^H NMR (500 MHz, DMSO_d6_) δ 9.21 (s, 1H), 4.95 (s, 2H), 4.14 (s, 2H), 3.62 (s, 3H), 3.35 (s, 3H), 2.65 (s, 3H); ^13^C NMR (126 MHz, DMSO_d6_) δ 169.5, 149.3, 147.2, 141.8, 141.0, 135.2, 128.6, 119.1, 65.4, 58.0, 53.0, 36.2, 14.0. Mass ESI [M+H]^+^ = 338.94.

#### Methyl 2-((8-bromo-9-methylthieno[3,2-*e*][1,2,4]triazolo[4,3-*c*]pyrimidin-3-yl)thio)acetate (8)

Off-white solid; yield: 65%; purity: >95%. ^1^H NMR (500 MHz, DMSO_d6_) δ 9.28 (s, 1H), 4.16 (s, 2H), 3.63 (s, 3H), 2.67 (s, 3H); ^13^C NMR (126 MHz, DMSO_d6_) δ 169.4, 151.2, 146.8, 141.4, 136.3, 132.1, 118.8, 112.7, 53.0, 36.1, 15.3. Mass ESI [M+H]^+^ = 372.87/374.86.

#### Methyl 2-((7,8,9-trimethyl-7*H*-pyrrolo[3,2-*e*][1,2,4]triazolo[4,3-*c*]pyrimidin-3-yl)thio)acetate (9)

White solid; yield: 60%; purity: >95%. ^1^H NMR (500 MHz, DMSO_d6_) δ 8.85 (s, 1H), 3.99 (s, 2H), 3.71 (s, 3H), 3.51 (s, 3H), 2.36 (d, *J* = 0.8 Hz, 3H), 2.30 (s, 3H); ^13^C NMR (126 MHz, DMSO_d6_) δ 169.5, 148.0, 138.7 138.6, 132.6, 131.0, 107.3, 102.1, 52.9, 36.3, 29.5, 10.6, 10.1. Mass ESI [M+H]^+^ = 306.00.

#### Methyl 2-((8,9-dimethylfuro[3,2-*e*][1,2,4]triazolo[4,3-*c*]pyrimidin-3-yl)thio)acetate (10)

Off-white solid; yield: 57%; purity: >95%. ^1^H NMR (500 MHz, DMSO_d6_) δ 9.11 (s, 1H), 4.11 (s, 2H), 3.61 (s, 3H), 2.47 (s, 3H), 2.38 (s, 3H); ^13^C NMR (126 MHz, DMSO_d6_) δ 169.5, 154.5, 150.9, 147.9, 140.3, 134.3, 110.9, 105.5, 53.0, 36.2, 11.9, 9.5. Mass ESI [M+H]^+^ = 292.97.

#### Methyl 2-((8,9-dimethoxy-[1,2,4]triazolo[4,3-*c*]quinazolin-3-yl)thio)acetate (11)

White solid; yield: 66%; purity: >95%. ^1^H NMR (500 MHz, DMSO_d6_) δ 9.03 (s, 1H), 7.71 (s, 1H), 7.44 (s, 1H), 4.06 (s, 2H), 3.93 (s, 3H), 3.89 (s, 3H), 3.53 (s, 3H); ^13^C NMR (126 MHz, DMSO_d6_) δ 169.5, 153.1, 151.0, 148.7, 141.5, 136.7, 134.6, 109.9, 102.7, 56.6, 56.5, 53.0, 36.2. Mass ESI [M+H]^+^ = 334.98.

#### Methyl (5,6-dimethyl-4-oxo-3,4-dihydrothieno[2,3-*d*]pyrimidine-3-carbonyl)glycinate (15)

Compound S1-0 (18 mg, 0.1 mmol) in DCM (2 mL) was added methyl 2-aminoacetate (9 mg, 0.1 mmol) and triethylamine (41 μL, 0.3 mmol). The mixture was cooled with ice and triphosgene (15 mg, 0.05 mmol) was added, stirred at room temperature for overnight. The resulting mixture was diluted with DCM and washed with water, saturated NaHCO_3_ and brine. The organic was dried with Na_2_SO_4_ and then concentrated under vacuum. The residue was purified by silica-gel column chromatography with 0–100 % ethyl acetate in hexanes to afford the final product. White solid; yield: 40%; purity: >95%. ^1^H NMR (500 MHz, DMSO_d6_) δ 9.97 (t, *J* = 5.7 Hz, 1H), 8.75 (s, 1H), 4.18 (d, *J* = 5.6 Hz, 2H), 3.70 (s, 3H), 2.42 (s, 3H), 2.41 (s, 3H); ^13^C NMR (126 MHz, DMSO_d6_) δ 169.8, 161.0, 158.7, 152.0, 142.9, 132.4, 130.1, 122.9, 52.5, 42.8, 13.6, 13.3. Mass ESI [M+H]^+^ = 296.01.

Using intermediate S4-0 and respective alkyl or benzyl bromide, compounds **16, 19, 21** and **22** were synthesized following General Procedure 4.

#### Ethyl 2-((8,9-dimethylthieno[3,2-*e*][1,2,4]triazolo[4,3-*c*]pyrimidin-3-yl)thio)acetate (16)

Off-white solid; yield: 67%; purity: >95%. ^1^H NMR (500 MHz, DMSO_d6_) δ 9.17 (s, 1H), 4.10 (s, 2H), 4.05 (q, *J* = 7.1 Hz, 2H), 2.64 (s, 3H), 2.55 (s, 2H), 1.09 (t, *J* = 7.1 Hz, 3H); ^13^C NMR (126 MHz, DMSO_d6_) δ 168.9, 149.0, 147.7, 140.7, 136.2, 134.8, 127.5, 119.4, 61.8, 36.4, 14.3, 13.7, 13.4. Mass ESI [M+H]^+^ = 322.95.

#### 3-(((8,9-dimethylthieno[3,2-*e*][1,2,4]triazolo[4,3-*c*]pyrimidin-3-yl)thio)methyl)dihydrofuran-2(3*H*)-one (19)

Light yellow solid; yield: 51%; purity: >95%. ^1^H NMR (500 MHz, DMSO_d6_) δ 9.20 (s, 1H), 4.60 (t, *J* = 9.0 Hz, 1H), 4.34 (td, *J* = 8.6, 3.6 Hz, 1H), 4.25 (td, J = 8.6, 7.0 Hz, 1H), 2.77 – 2.70 (m, 1H), 2.65 (s, 3H), 2.55 (s, 3H), 2.48 – 2.42 (m, 1H); ^13^C NMR (126 MHz, DMSO_d6_) δ 174.8, 149.3, 148.0, 138.8, 136.4, 134.8, 127.5, 119.4, 67.2, 44.7, 29.7, 13.7, 13.4. Mass ESI [M+H]^+^ = 320.93.

#### 3-((3,5-difluoro-4-methoxybenzyl)thio)-8,9-dimethylthieno[3,2-e][1,2,4]triazolo[4,3-c]pyrimidine (21)

Light yellow solid; yield: 53%; purity: >95%. ^1^H NMR (500 MHz, DMSO_d6_) δ 9.00 (s, 1H), 7.12 – 7.07 (m, 2H), 4.34 (s, 2H), 3.84 (s, 3H), 2.63 (s, 3H), 2.54 (s, 3H); ^13^C NMR (126 MHz, DMSO_d6_) δ 155.1 (dd, *J* = 246.9, 6.7 Hz), 149.1, 147.8, 140.4, 136.1, 135.4 (t, *J* = 14.1 Hz), 134.5, 133.9 (t, *J* = 9.3 Hz), 127.5, 119.4, 113.7 (dd, *J* = 17.8, 6.2 Hz), 62.2, 37.4, 13.7, 13.4. Mass ESI [M+H]^+^ =393.01.

#### Methyl 3-((8,9-dimethylthieno[3,2-*e*][1,2,4]triazolo[4,3-*c*]pyrimidin-3-yl)thio)propanoate (22)

Light yellow solid; yield: 63%; purity: >95%. ^1^H NMR (500 MHz, DMSO_d6_) δ 9.11 (s, 1H), 3.56 (s, 3H), 3.35 (t, *J* = 6.9 Hz, 2H), 2.80 (t, *J* = 6.8 Hz, 2H), 2.64 (s, 3H), 2.54 (s, 3H); ^13^C NMR (126 MHz, DMSO_d6_) δ 172.0, 149.0, 147.8, 140.9, 136.0, 134.6, 127.5, 119.5, 52.0, 34.5, 29.4, 13.7, 13.4. Mass ESI [M+H]^+^ = 322.96.

#### Methyl 2-(8,9-dimethylthieno[3,2-*e*][1,2,4]triazolo[4,3-*c*]pyrimidin-3-yl)acetate (23)

Compound S3-0 (100 mg, 0.51 mmol) in dimethyl malonate (2 mL) was heated at 180 °C for 45 min under microwave. After cooling, solvent was removed under vacuum and the residue was purified by silica-gel column chromatography with 0–50 % ethyl acetate in hexanes to afford the final product. White solid; yield: 60%; purity: >95%. ^1^H NMR (500 MHz, DMSO_d6_) δ 9.57 (s, 1H), 4.10 (s, 2H), 3.69 (s, 3H), 2.59 (s, 3H), 2.53 (s, 3H); ^13^C NMR (126 MHz, DMSO_d6_) δ 169.6, 161.6, 152.3, 149.2, 137.2, 136.1, 126.8, 121.0, 52.6, 35.0, 13.8, 13.1. Mass ESI [M+H]^+^ = 277.00.

#### Methyl 2-((8,9-dimethylthieno[3,2-*e*][1,2,4]triazolo[4,3-*c*]pyrimidin-3-yl)oxy)acetate (24)

Compound S3-0 (50 mg, 0.25 mmol) in acetonitrile (2 mL) was added 1,1’-carbonyldiimidazole (40 mg, 0.25 mmol). The mixture was heated at 90 °C for 1 h. After cooling, the mixture was poured onto ice and then extracted with ethyl acetate. The organic was washed with brine, dried with Na_2_SO_4_ and concentrated. The residue was purified by silica-gel column chromatography with 0–10 % MeOH in DCM to give compound S6, 30 mg (53% yield). Mass ESI [M+H]^+^ = 221.02.

Compound S6 (30 mg, 0.13 mmol) was dissolved in anhydrous DMF (1 mL) and cooled with ice. NaH (60% in oil, 6 mg, 0.15 mmol) was added and the mixture was stirred for 15 min. Methyl bromoacetate (18 μL, 0.19 mmol) was added and the mixture was stirred at room temperature for 30 min. The mixture was poured onto ice and then extracted with ethyl acetate. The organic was washed with brine, dried with Na_2_SO_4_ and concentrated. The residue was purified by silica-gel column chromatography with 0–50 % ethyl acetate in hexanes to afford the final compound. White solid; yield: 45%; purity: 92.8%. ^1^H NMR (500 MHz, DMSO_d6_) δ 8.65 (s, 1H), 4.87 (s, 2H), 3.73 (s, 3H), 2.46 (s, 3H), 2.43 (s, 3H); ^13^C NMR (126 MHz, DMSO_d6_) δ 168.4, 150.4, 147.8, 138.1, 135.6, 135.0, 127.4, 117.8, 53.0, 47.2, 13.4, 13.1. Mass ESI [M+H]^+^ =292.99.

#### Methyl 3-(8,9-dimethylthieno[3,2-*e*][1,2,4]triazolo[4,3-*c*]pyrimidin-3-yl)propanoate (25)

Compound S3-0 (100 mg, 0.25 mmol) in DMF (2 mL) was added succinic anhydride (37 mg, 0.37 mmol). The mixture was heated at 80 °C for overnight. After cooling, the mixture was poured onto ice and then extracted with ethyl acetate. The organic was washed with brine, dried with Na_2_SO_4_ and concentrated. The residue was purified by silica-gel column chromatography with 0–5 % MeOH in DCM to give compound S7, 20 mg (14% yield). ^1^H NMR (400 MHz, DMSO_d6_) δ 9.54 (s, 1H), 3.15 (t, *J* = 7.2 Hz, 2H), 2.83 (t, *J* =7.2 Hz, 2H), 2.61 (s, 3H), 2.54 (s, 3H). Mass ESI [M+H]^+^ = 277.06.

Compound S7 (20 mg, 0.07 mmol) was dissolved in methanol (1 mL) and added one drop conc. H_2_SO_4_. The mixture was refluxed at 80 °C for 2 h. Methanol was removed under vacuum. The residue was dissolved in ethyl acetate, washed with saturated NaHCO_3_ and brine, dried with Na_2_SO_4_ and concentrated. The crude product was purified by silica-gel column chromatography with 0–20 % ethyl acetate in hexanes to afford the final compound. White solid; yield: 71%; purity: >95%. ^1^H NMR (500 MHz, DMSO_d6_) δ 9.52 (s, 1H), 3.63 (s, 3H), 3.19 (t, *J* = 7.2 Hz, 3H), 2.91 (t, *J* = 7.2 Hz, 3H), 2.59 (s, 3H), 2.53 (s, 3H); ^13^C NMR (126 MHz, DMSO_d6_) δ 172.9, 166.7, 152.2, 149.1, 137.2, 135.8, 126.8, 121.0, 51.9, 31.4, 24.1, 13.7, 13.1. Mass ESI [M+H]^+^ = 290.99.

#### Methyl 2-((8,9-dimethylthieno[3,2-*e*][1,2,4]triazolo[4,3-*c*]pyrimidin-3-yl)sulfinyl)acetate (26)

Compound **BDW568** (20 mg, 0.06 mmol) was dissolved in DCM (1 mL) and cooled with ice. mCPBA (10 mg, 0.06 mmol) was added and the mixture was stirred at room temperature for 2 h. The resulting mixture was diluted with DCM and washed with saturated NaHCO_3_ and brine. The organic was dried with Na_2_SO_4_ and then concentrated under vacuum. The residue was purified by silica-gel column chromatography with 0–100 % ethyl acetate in hexanes to afford the final product. Off-white solid; yield: 32%; purity: >95%. ^1^H NMR (500 MHz, DMSO_d6_) δ 9.38 (s, 1H), 5.22 (s, 2H), 3.62 (s, 3H), 2.71 (s, 3H), 2.60 (s, 3H); ^13^C NMR (126 MHz, DMSO_d6_) δ 163.6, 150.4, 148.1, 143.6, 137.7, 134.1, 127.8, 119.5, 60.4, 53.5, 13.8, 13.5. Mass ESI [M+H]^+^ = 340.95.

#### THP-1 ISRE reporter assay

THP-1 ISRE reporter cells (InvivoGen, thpd-nfis) were seeded in 384-well plates with 5,000 cells per well in a total of 30 μL of full medium containing RPMI 1640 (Gibco, 31-800-022), 10% fetal bovine serum (FBS) (HyClone, SH3039603HI), 2 g/L sodium bicarbonate, 1 mM sodium pyruvate, 10 mM HEPES buffer (pH 7.3), 1× Antibiotic-Antimycotic (Gibco, 15-240-062), and 1 mM β-mercaptoethanol. The cells were treated with a 1:2 serial dilution of BDW568 (or analogs) at 12 concentrations in quadruplicates. Compounds were incubated with the reporter cells for 48 h before 10 μL of 0.5 × QUANTI-Luc (InvivoGen, rep-qlc2) was added to each well. The plates were measured immediately using a luminescence plate reader. EC_50_’s were determined with GraphPad prism 9.5.1 nonlinear regression analysis, and shown as mean ± s.d..

#### Lentivirus construction, production and purification

The lentiviral vector FG12-EF1p-EGFRt-p2A-STING^230A^ was constructed by cloning STING^230A^ variant into a FG12 expression vector. The vector also contains EF1α promoter sequence that expresses the α subunit of eukaryotic elongation factor 1 along with truncated inactive epidermal growth factor receptor (EGFRt) that can be used as flow-cytometric marker and a self-cleaving peptide 2A (p2A). The lentivirus vector was produced in 293FT cells using the calcium phosphate transfection protocol. Briefly, DNA mix was prepared by adding above mentioned STING^230A^ plasmid with the third-generation lentiviral packaging plasmid pMDLg along with Rev and VSVG envelope expressing plasmid (Azenta Life sciences, USA). The DNA was mixed with 2 M CaCl_2_ in cell-grade water followed by addition of 2× HEPES and 2× HBS. The mixture was incubated at room temperature for 25 min and added to 293FT cells cultured in IMDM complete media (IMDM media, 10% FBS, 1% PBS, 1% Glutamax) and 10mM chloroquine. After 6–8 h of incubation, the transfection media was aspirated and fresh 2% Opti-MEM compete media (Opti-MEM media with 2% FBS, 1% Pen-Strep, 1% Glutamax) was added followed by 2 days of incubation. Supernatant was collected from transfected 293FT cells 48 h following transfection, filtered using a 0.45 μm sterile filter, and concentrated by ultracentrifugation using a Beckman SW32 rotor at 30,000 rpm at 4°C. Medium was aspirated and pellet was resuspended with PBS and stored at –80°C.

#### Primary macrophage purification and differentiation

Healthy human PBMCs were obtained from the Department of Virology, UCLA AIDS institute. Monocytes were magnetically sorted using human CD14^+^ microbeads (Miltenyi Biotec, USA) according to manufactures protocol. Briefly, PBMCs were incubated with CD14^+^ microbeads and passed through LS columns placed in a magnetic field with multiple rounds of washing using MACS buffer. CD14^+^ cells were differentiated into macrophages in the presence of RPMI complete media (RPMI 1640 media, 10%FBS, 1% PenStrep) and macrophage colony-stimulating factor (M-CSF) growth factor at a concentration of 10 ng/ml for 5 days. Macrophages were transduced with FG12-EF1p-EGFRt-p2A-STING^230A^ lentivirus or mock transduced at MOI:10 overnight and washed with fresh media afterwards. Cells were cultured for 2 more days in the presence of RPMI complete media and M-CSF. Three days after transduction, untransduced and transduced macrophages were stimulated with DMSO (control) and BDW568 at a concentration of 50 μM for 6 h. Cells were harvested using accutase for RNA extraction.

#### RT-qPCR

THP-1 and primary macrophage RNA was extracted using RNeasy kit (Qiagen), followed by cDNA synthesis using the High-Capacity cDNA Reverse Transcription Kit (Thermo Fisher Scientific). RT-qPCR was performed using the TaqMan Gene Expression Assays (Thermo Fisher Scientific), targeting human HPRT1 (Hs01003267_m1), MX1 (Hs00895608_m1) and OAS1 (Hs00973635_m1). To determine the relative mRNA expression, MX1 and OAS1 gene expression was normalized to the HPRT1 expression level.

#### Genotyping for STING residue 230

The gene fragment containing the residue 230 of STING was amplified by PCR using Phusion high-fidelity PCR kit (ThermoFisher Scientific, USA). The PCR reaction was set up using template genomic DNA, Phusion DNA polymerase, STING230-GT forward primer 5’-GGCCCGGATTCGAACTTACA and STING230-GT reverse primer 5’-CCTGCACCCACATGAATCCT. Following PCR, the generated DNA fragment was purified from agarose gel and subcloned into a pCR™Blunt II-TOPO™ vector using Zero Blunt TOPO PCR cloning kit (ThermoFisher Scientific, USA). Afterwards, colonies were picked, and the plasmid DNA were isolated using QIAprep Spin Miniprep kit (Qiagen) and Sanger sequenced (Laragen, Inc. CA).

#### Supporting Information

Data collection and refinement statistics for STING^A230^–ligand complexes; ^1^H, ^13^C NMR Spectra; purity analysis of active compounds.

## AUTHOR INFORMATION

### Author Contributions

Z.T., M.S., and J.W. performed the chemical synthesis. Y.L., D.S., and P.L. expressed the STING^A230^ protein, performed crystallography, and analyzed the structural data. J.Z., Y.L., and D.K.J. visualized the structures. J.Z., S.T., N.T., and J.W. performed biological evaluation in THP-1 cells and analyzed the data. S.T., N.T., and A.Z. performed STING stimulation experiments in primary cells. J.W. wrote the manuscript with input from all the authors.

## ACKNOWLEDGMENT

The research described in the manuscript was funded by the National Institutes of Health, grant number P20GM103638 (J.W.), R01AI145287 (P.L.), R01DA052841 (A.Z.), R01AI172727 (A.Z.), and R21AI155117 (A.Z.), and the University of Kansas Research Grant Opportunity (J.W.). The work at UCLA was also supported by the UCLA AIDS Institute, the James B. Pendleton Charitable Trust, and the McCarthy Family Foundation. We thank Wenshe Liu at Texas A&M University for discussions.

## Notes

### Competing Interest Statement

The authors have declared no competing interest.

### Summary of Updates

Typos in Figures 3A & 3B (T263 was mislabeled as T163 in the first version) and the Method section "Crystallization and structure determination" were corrected. The section title "Crystal structure of STING-A230 CTD-BDW-OH complex" was also corrected.

