## Supplementary Information for "Structural and Biological Evaluations of a Non-Nucleoside STING Agonist Specific for Human STING^A230^ Variants"

**Table S1**. Data collection and refinement statistics for STING$^{A230}$ CTD–ligand complexes

| | **STING-cGAMP** | **STING-BDW568** |
|---|---|---|
| **Data collection** | | |
| Space group | C 2 | C 2 |
| Molecules per ASU | 1 STING, 0.5 cGAMP | 1 STING, 1 ligand |
| Cell dimensions | | |
| *a*, *b*, *c* (Å) | 88.33, 79.23, 36.59 | 90.34, 77.11, 36.16 |
| $\alpha$, $\beta$, $\gamma$ (°) | 90.0, 96.13, 90.0 | 90.0, 97.96, 90.00 |
| Resolution (Å) | 2.01 (2.06 to 2.01)* | 1.95 (2.00 to 1.95)* |
| $R_{merge}$ | 17.7% (66.5%) | 11.3% (100.5%) |
| $R_{pim}$ | 10.8% (42.1%) | 7.2% (78.1%) |
| CC(1/2) (%) | 91.6 (62.4) | 92.9 (37.6) |
| Unique reflections | 16799 (1224) | 17718 (1112) |
| $I / \sigma I$ | 8.2 (1.9) | 9.8 (0.8) |
| Completeness (%) | 99.9 (99.1) | 98.7 (87.8) |
| Redundancy | 3.7 (3.4) | 3.6 (2.7) |
| **Refinement** | | |
| Resolution (Å) | 39.6 to 2.01 | 58.4 to 1.95 |
| No. reflections ($F > 0$) | 16789 | 17705 |
| $R_{work}$ / $R_{free}$ | 17.8% /22.0% | 19.8% /23.0% |
| No. atoms | | |
| Protein | 1432 | 1447 |
| Ligand/Water | 45/165 | 19/86 |
| *B*-factors (Å$^2$) | | |
| Protein | 39.0 | 56.6 |
| Water | 22.0/43.0 | 40.4/52.1 |
| R.m.s. deviations | | |
| Bond lengths (Å) | 0.006 | 0.007 |
| Bond angles (°) | 0.905 | 0.857 |
| Ramachandran plot favored (%) | 98.2 | 97.1 |
| Ramachandran plot outlier (%) | 0.0 | 0.0 |

* One crystal was used to collect each of the dataset.

*Values in parentheses are for highest-resolution shell.

Updated for refine_24 and refine_46, 6/15/23, deposited in PDB

**$^{1}H$ and $^{13}C$ NMR spectra**

Compound 1

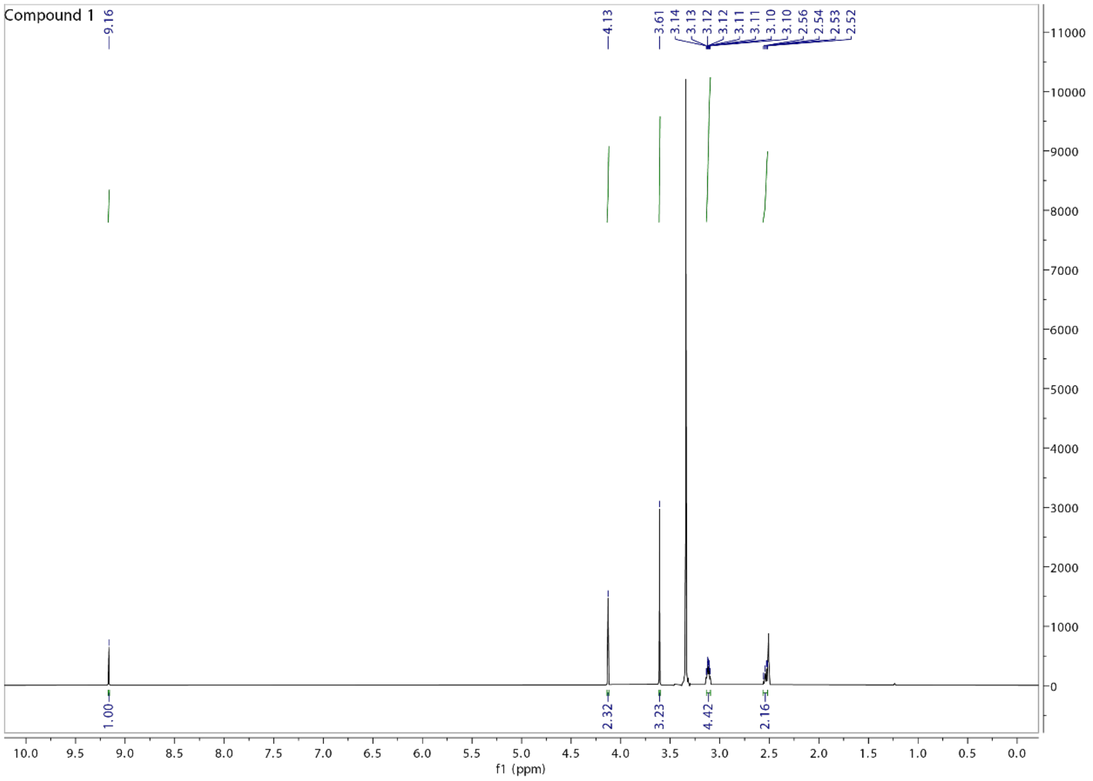

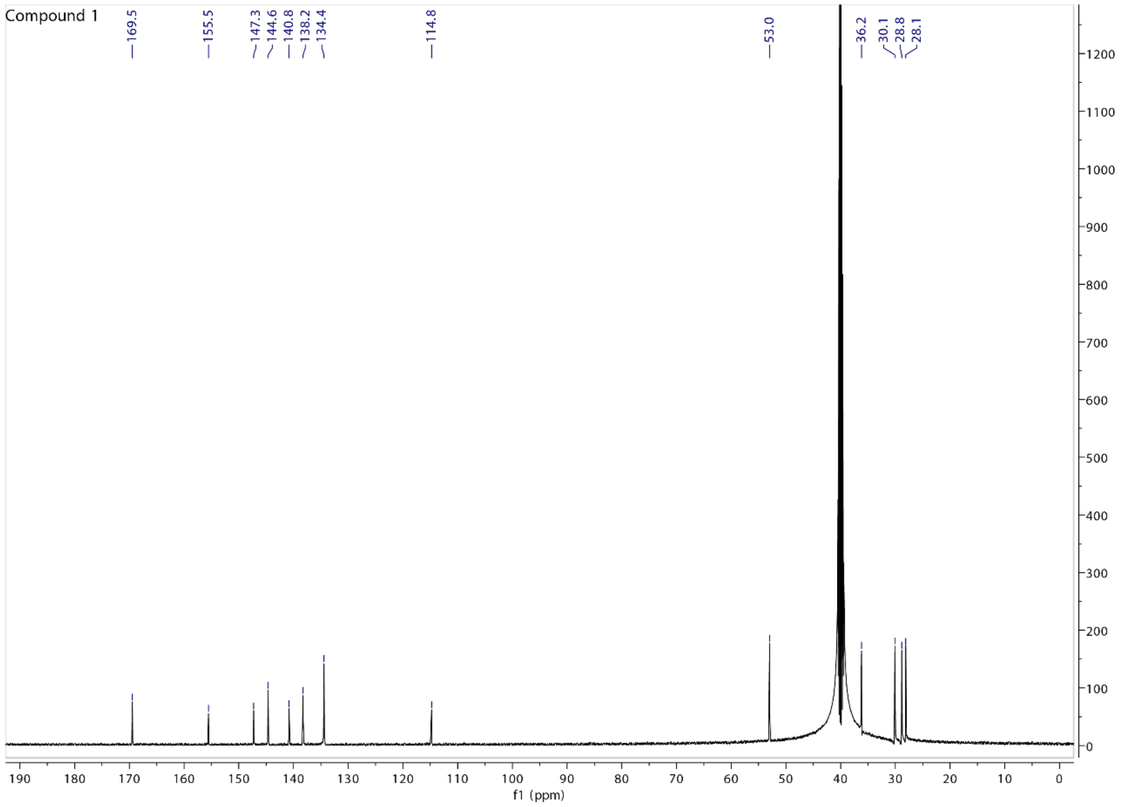

Compound 2

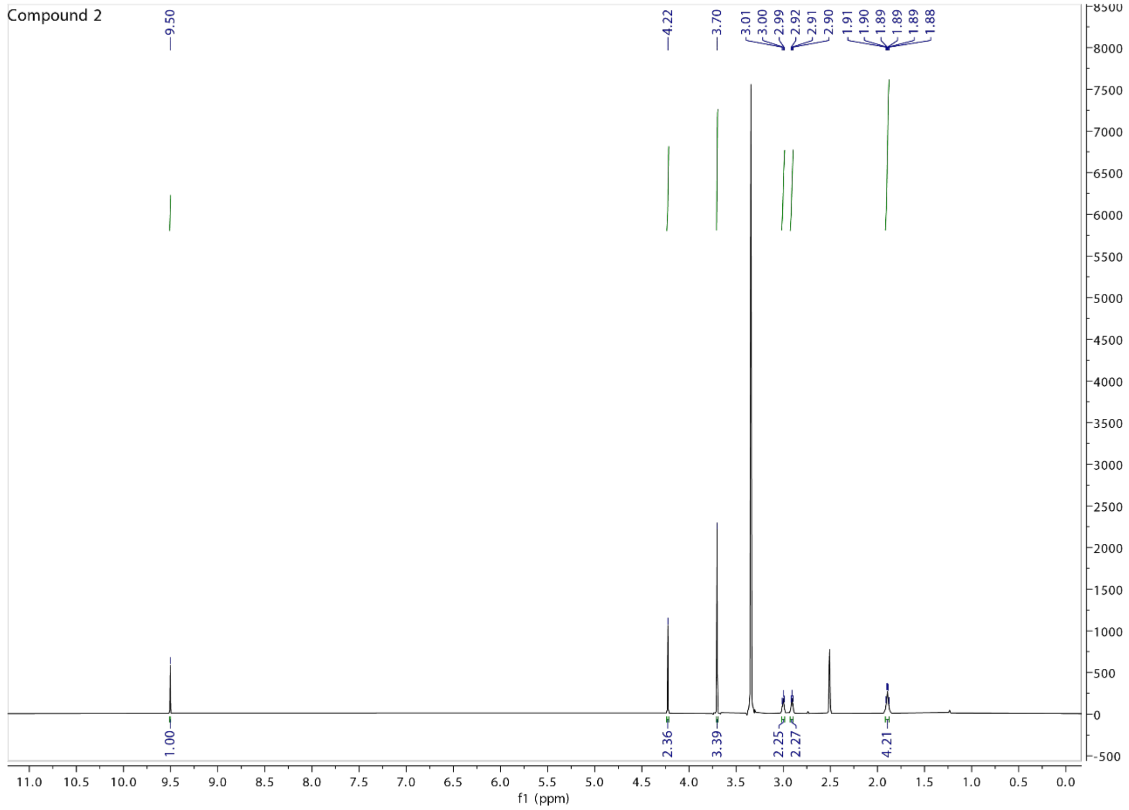
Compound 2
9.50
4.22
3.70
3.01
3.00
2.99
2.92
2.91
2.90
1.91
1.90
1.89
1.89
1.89
1.88
1.00
2.36
3.39
2.25
2.27
4.21
f1 (ppm)

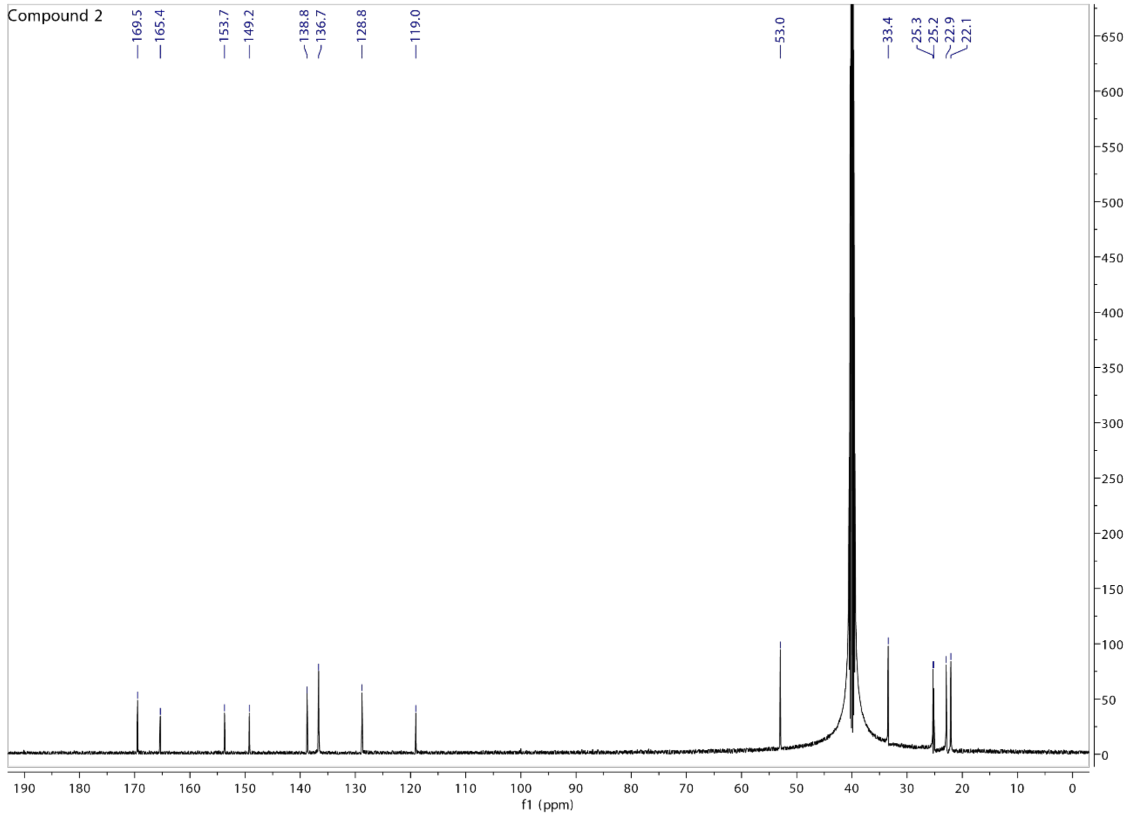
Compound 2
169.5
165.4
153.7
149.2
138.8
136.7
128.8
119.0
53.0
33.4
25.3
25.2
22.9
22.1
f1 (ppm)

Compound 3

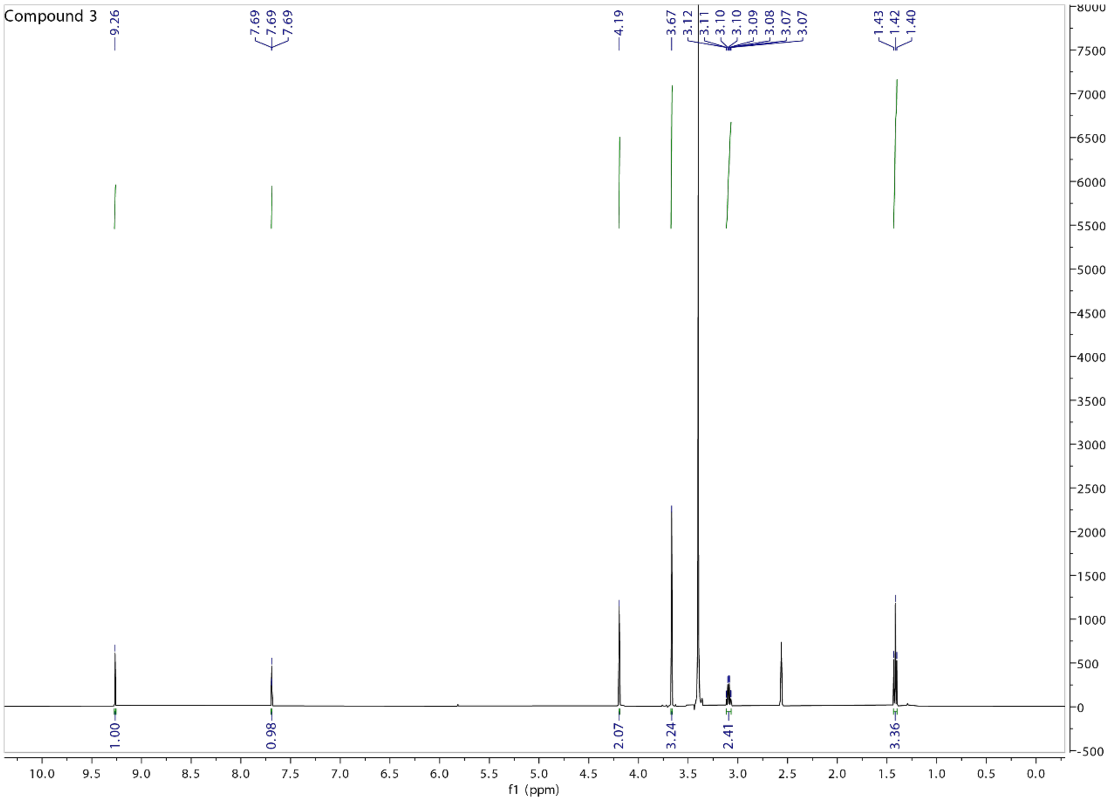

Compound 3
9.26
7.69
7.69
7.69
4.19
3.67
3.12
3.11
3.10
3.10
3.09
3.08
3.07
3.07
1.43
1.42
1.40
1.00
0.98
2.07
3.24
2.41
3.36
f1 (ppm)

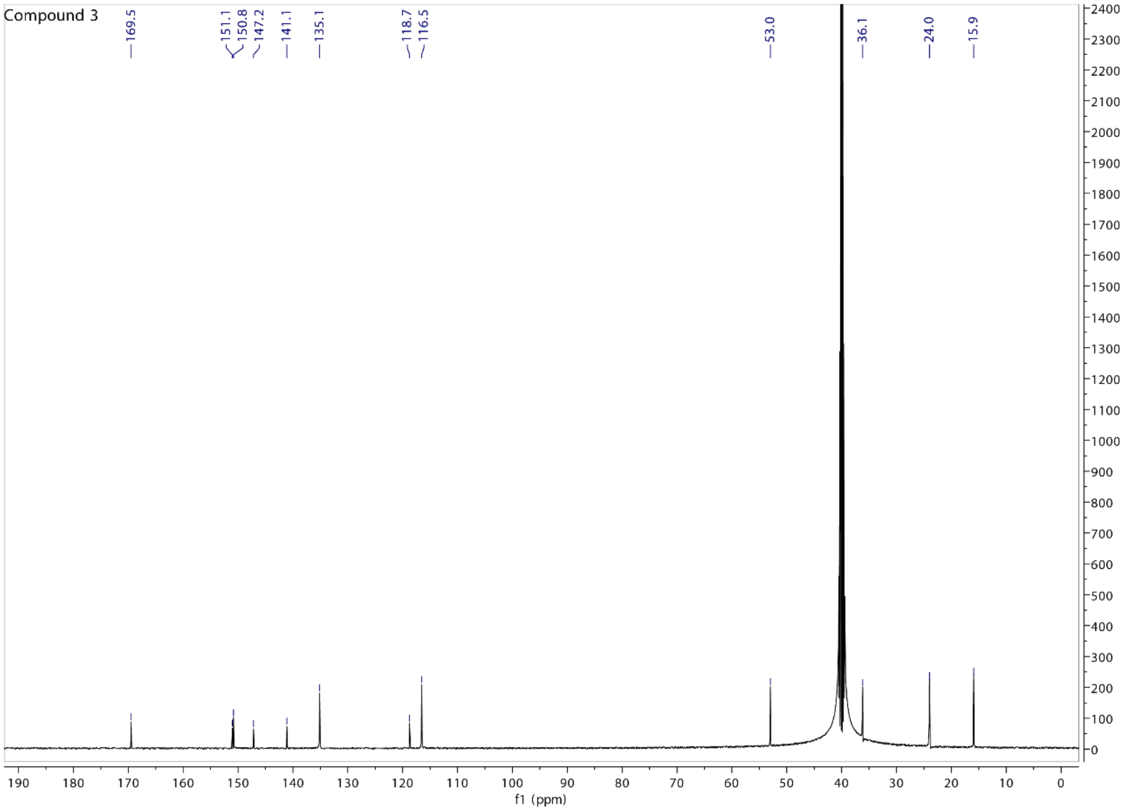

Compound 3
169.5
151.1
150.8
147.2
141.1
135.1
118.7
116.5
53.0
36.1
24.0
15.9
f1 (ppm)

Compound 4

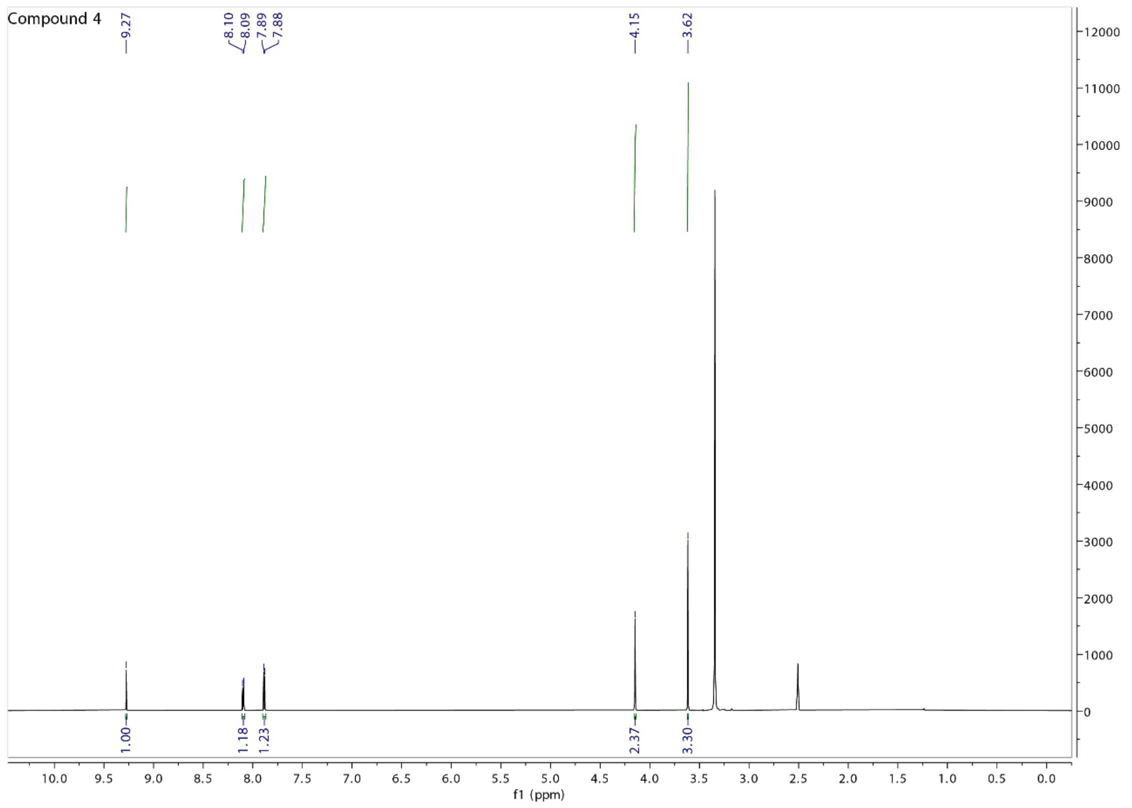
Compound 4
9.27
8.10
8.09
7.89
7.88
4.15
3.62
1.00
1.18
1.23
2.37
3.30
f1 (ppm)

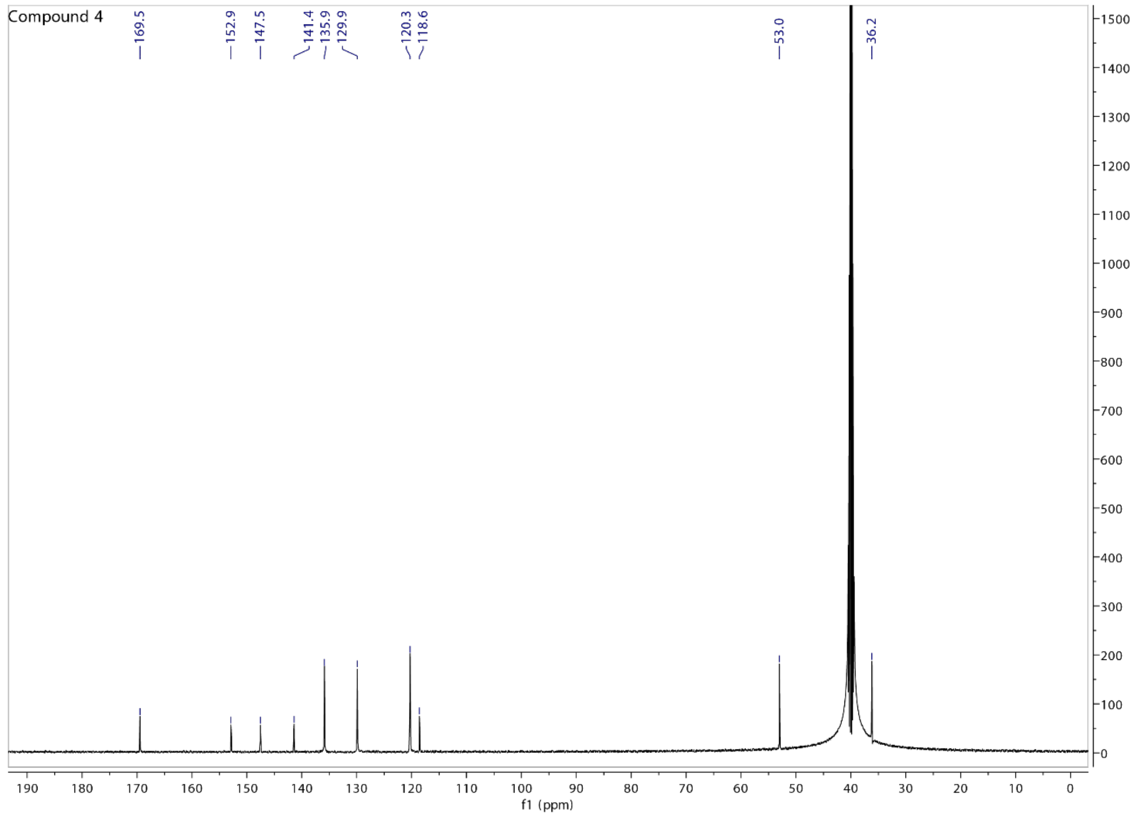
Compound 4
169.5
152.9
147.5
141.4
135.9
129.9
120.3
118.6
53.0
36.2
f1 (ppm)

#### Compound 5

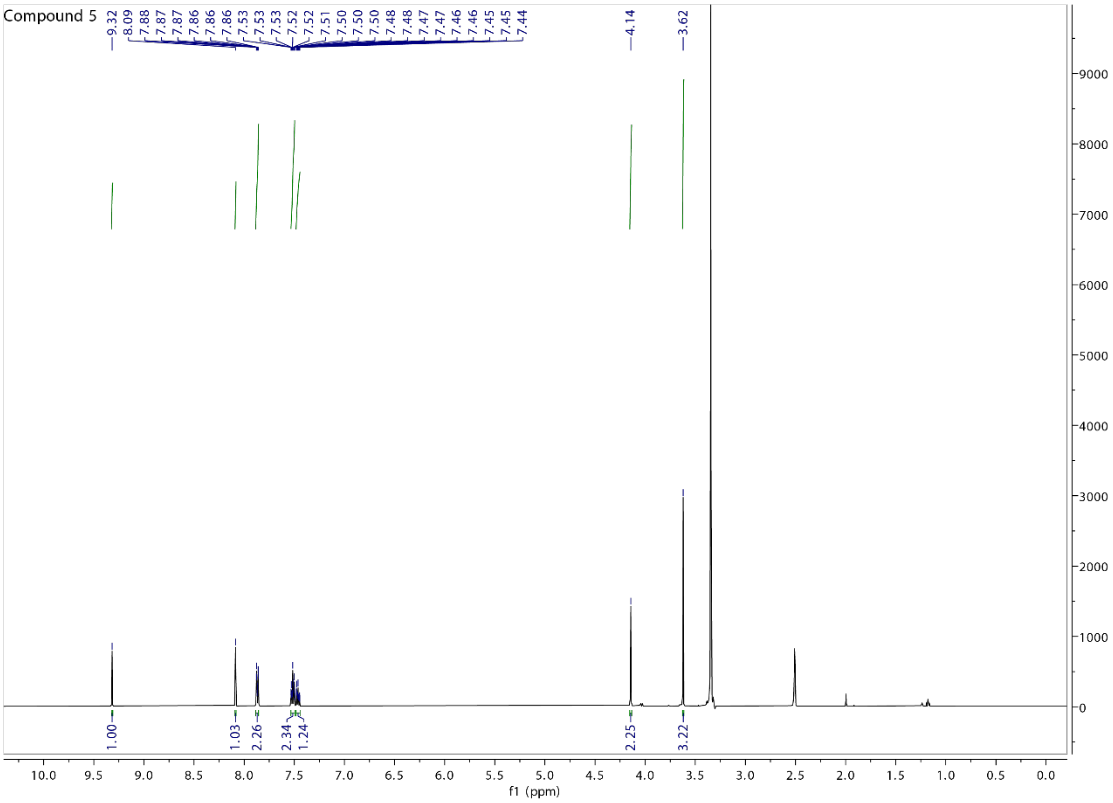
Compound 5
9.32
8.09
7.88
7.87
7.87
7.86
7.86
7.53
7.53
7.53
7.52
7.52
7.51
7.50
7.50
7.50
7.48
7.48
7.47
7.47
7.46
7.46
7.45
7.45
7.44
4.14
3.62
1.00
1.03
2.26
2.34
1.24
2.25
3.22
f1 (ppm)

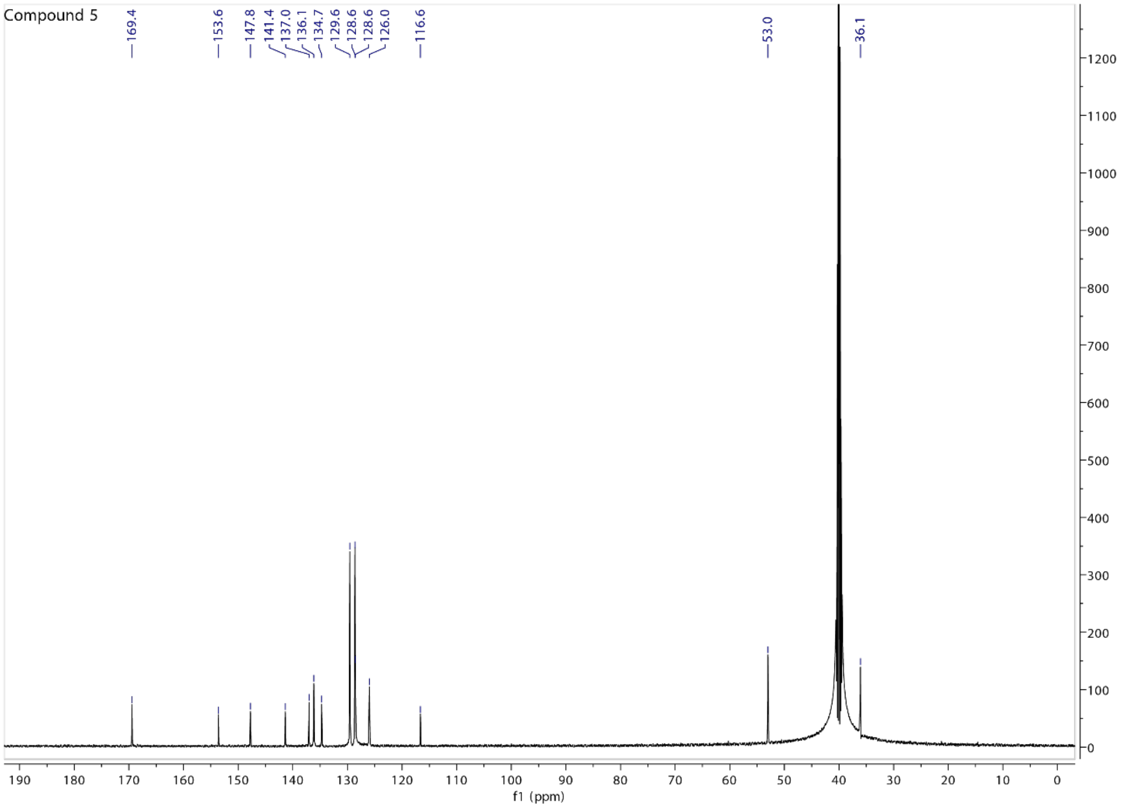
Compound 5
169.4
153.6
147.8
141.4
137.0
136.1
134.7
129.6
128.6
128.6
126.0
116.6
53.0
36.1
f1 (ppm)

Compound 6

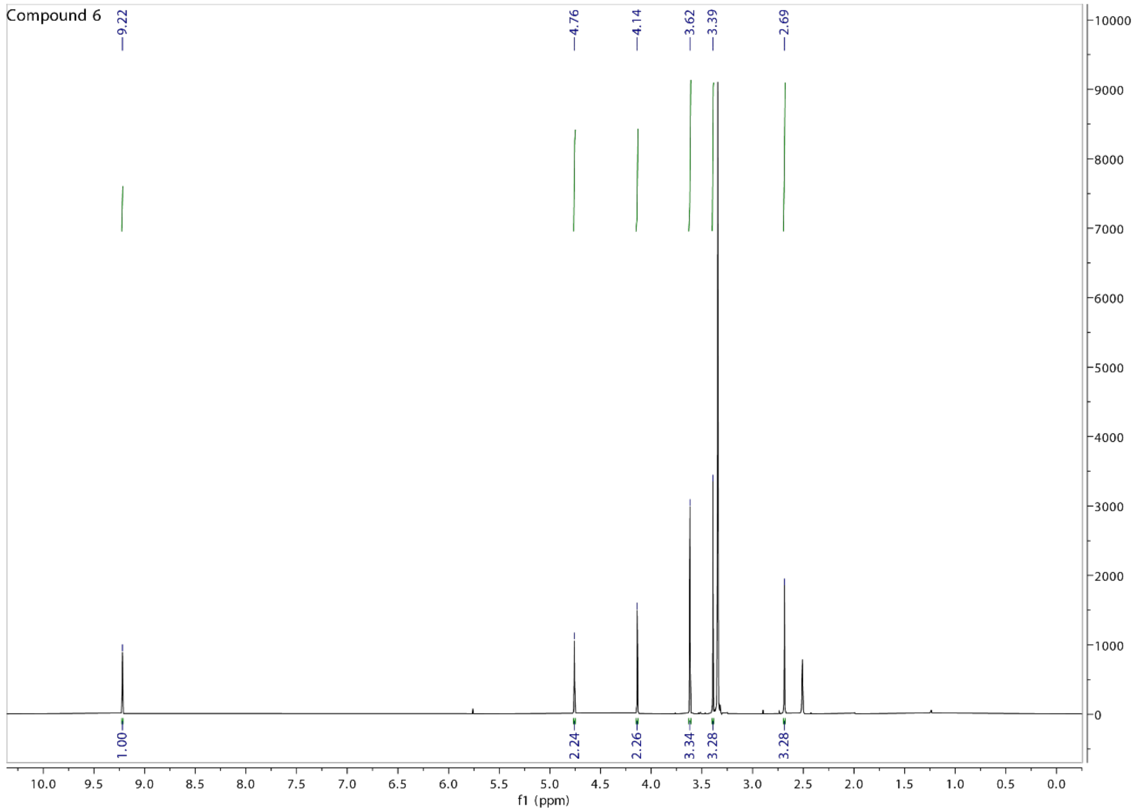
Compound 6
9.22
4.76
4.14
3.62
3.39
2.69
1.00
2.24
2.26
3.34
3.28
3.28
f1 (ppm)

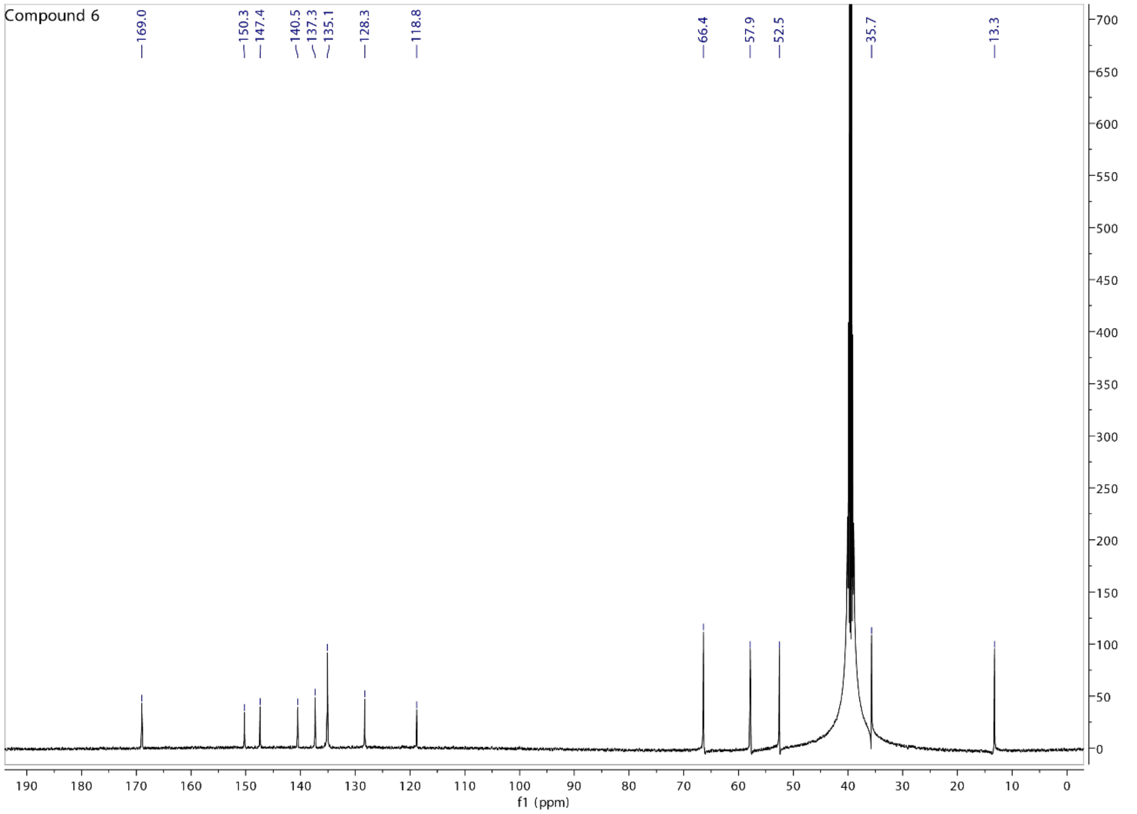
Compound 6
169.0
150.3
147.4
140.5
137.3
135.1
128.3
118.8
66.4
57.9
52.5
35.7
13.3
f1 (ppm)

Compound 7

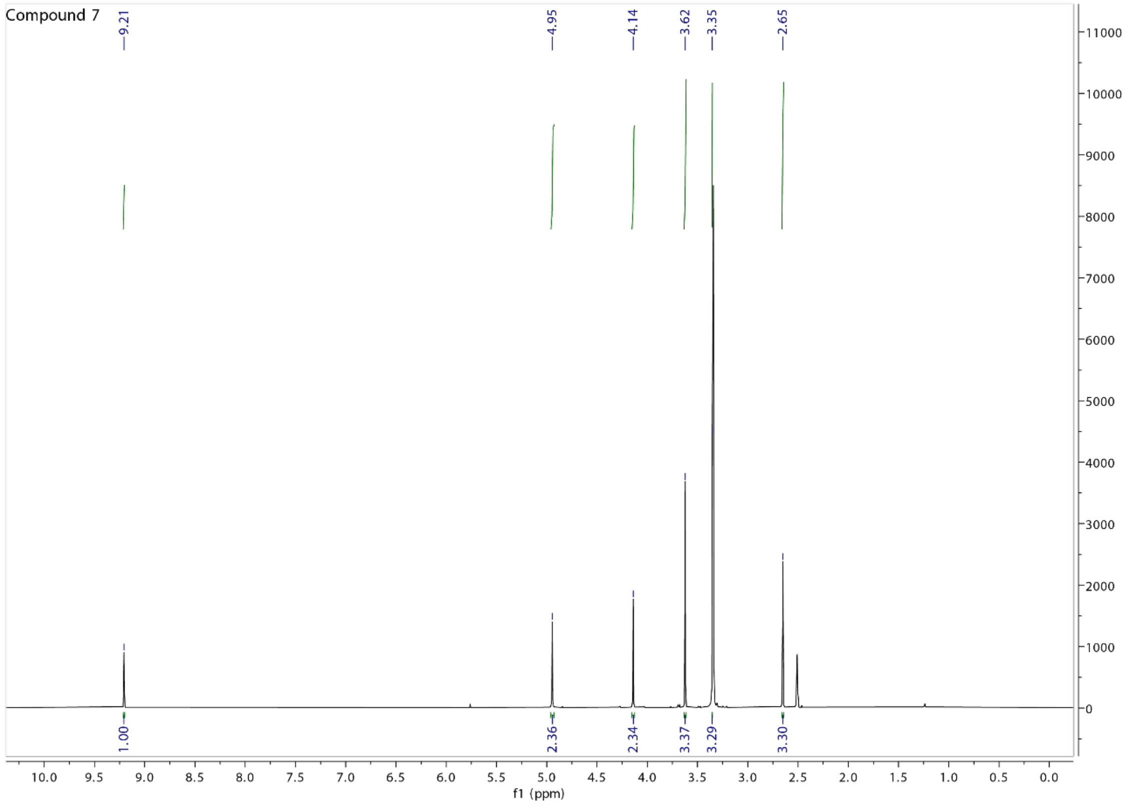
Compound 7
9.21
4.95
4.14
3.62
3.35
2.65
1.00
2.36
2.34
3.37
3.29
3.30
f1 (ppm)

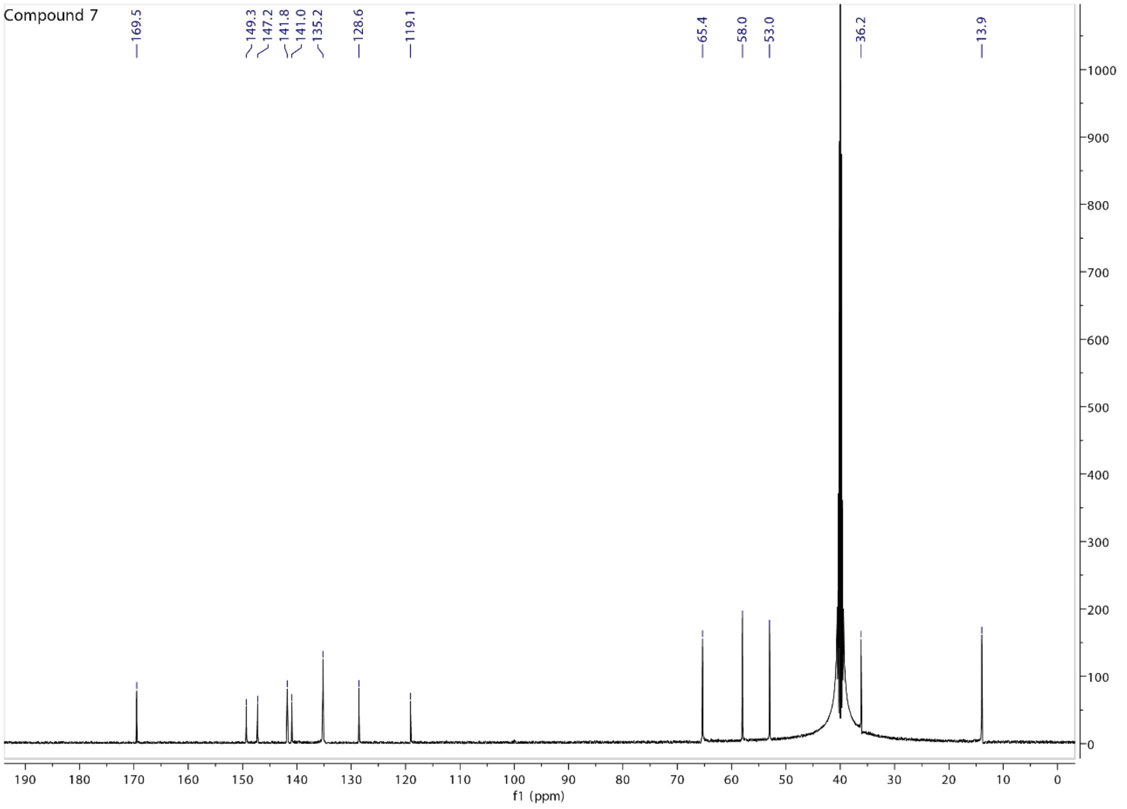
Compound 7
169.5
149.3
147.2
141.8
141.0
135.2
128.6
119.1
65.4
58.0
53.0
36.2
13.9
f1 (ppm)

### Compound 8

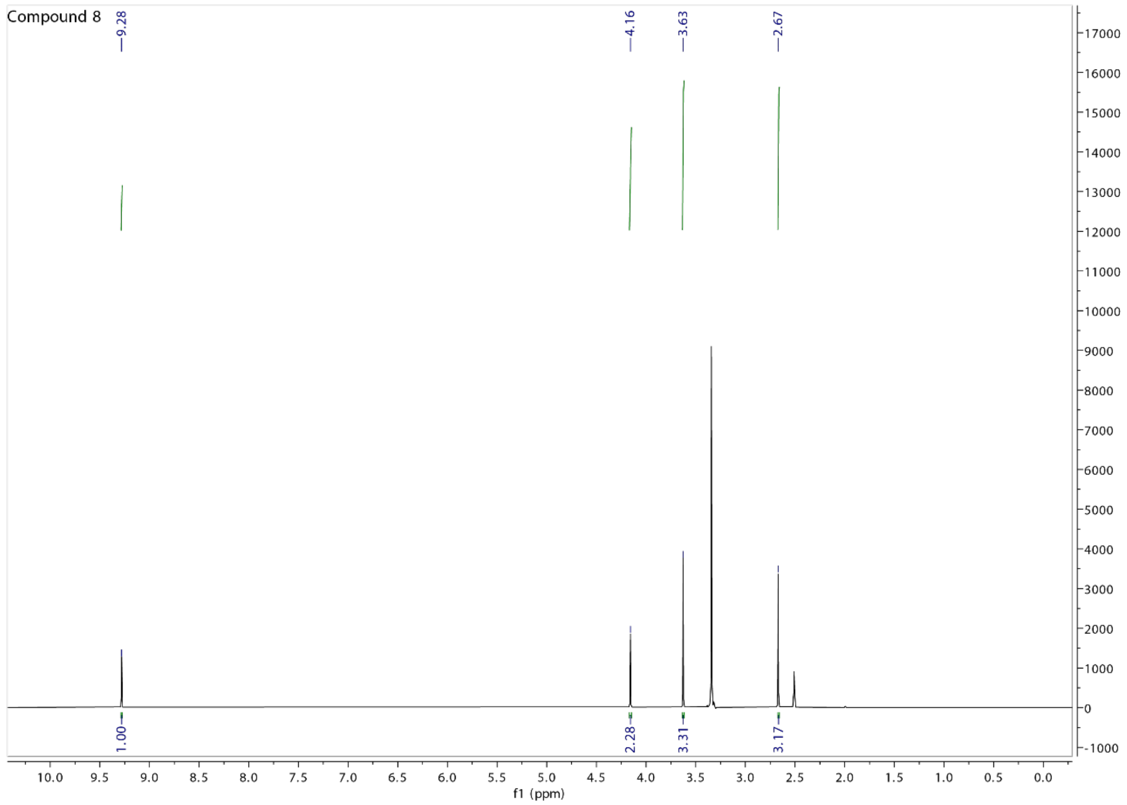
Compound 8
9.28
4.16
3.63
2.67
1.00
2.28
3.31
3.17
f1 (ppm)

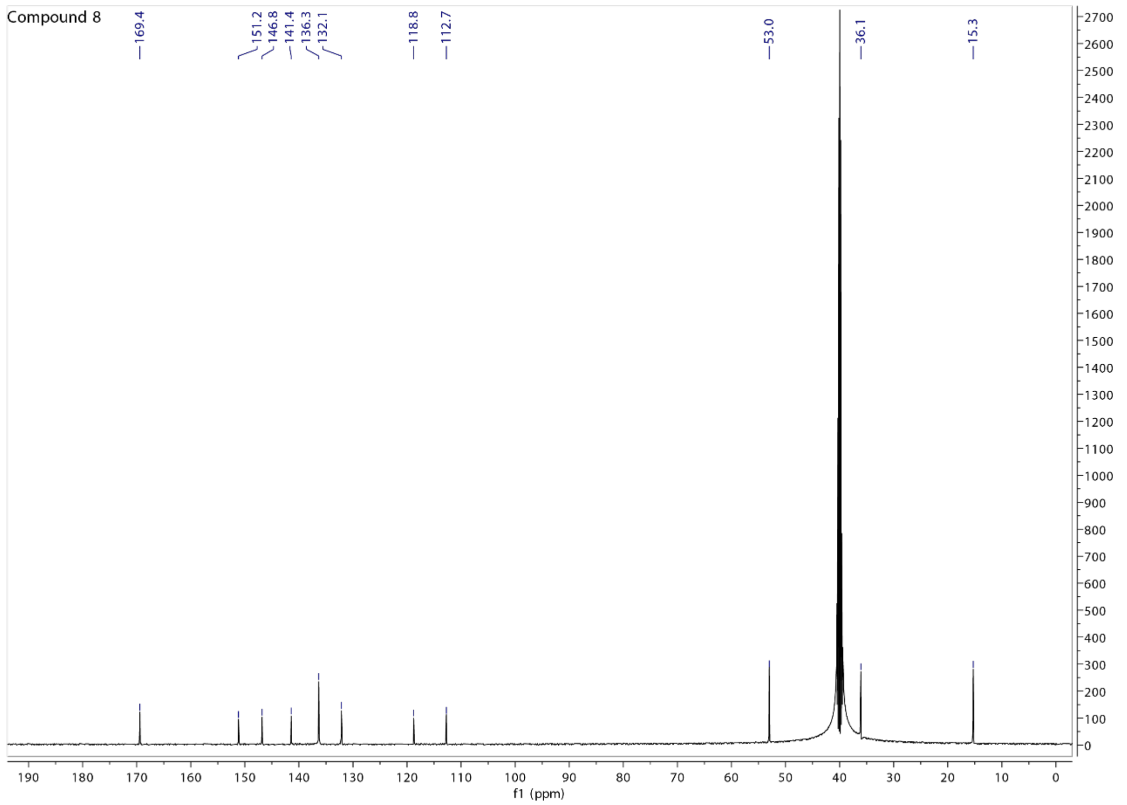
Compound 8
169.4
151.2
146.8
141.4
136.3
132.1
118.8
112.7
53.0
36.1
15.3
f1 (ppm)

Compound 9

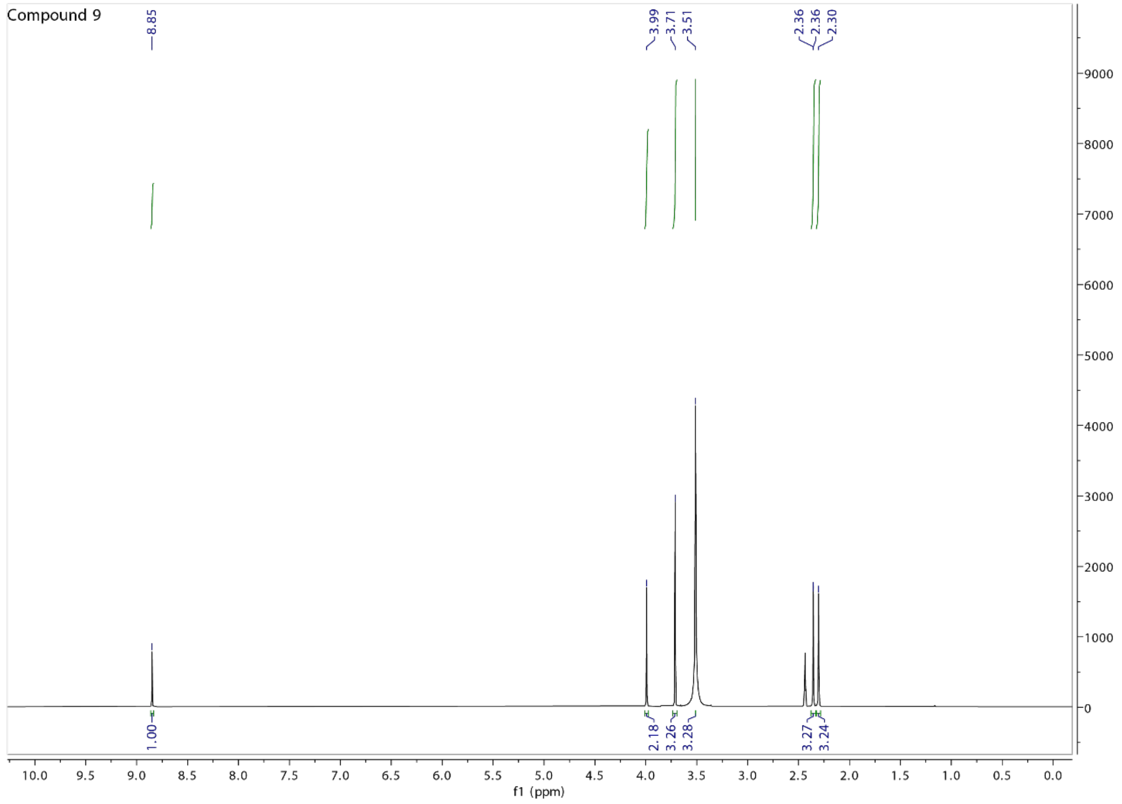
Compound 9
8.85
3.99
3.71
3.51
2.36
2.36
2.30
1.00
2.18
3.26
3.28
3.27
3.24
f1 (ppm)

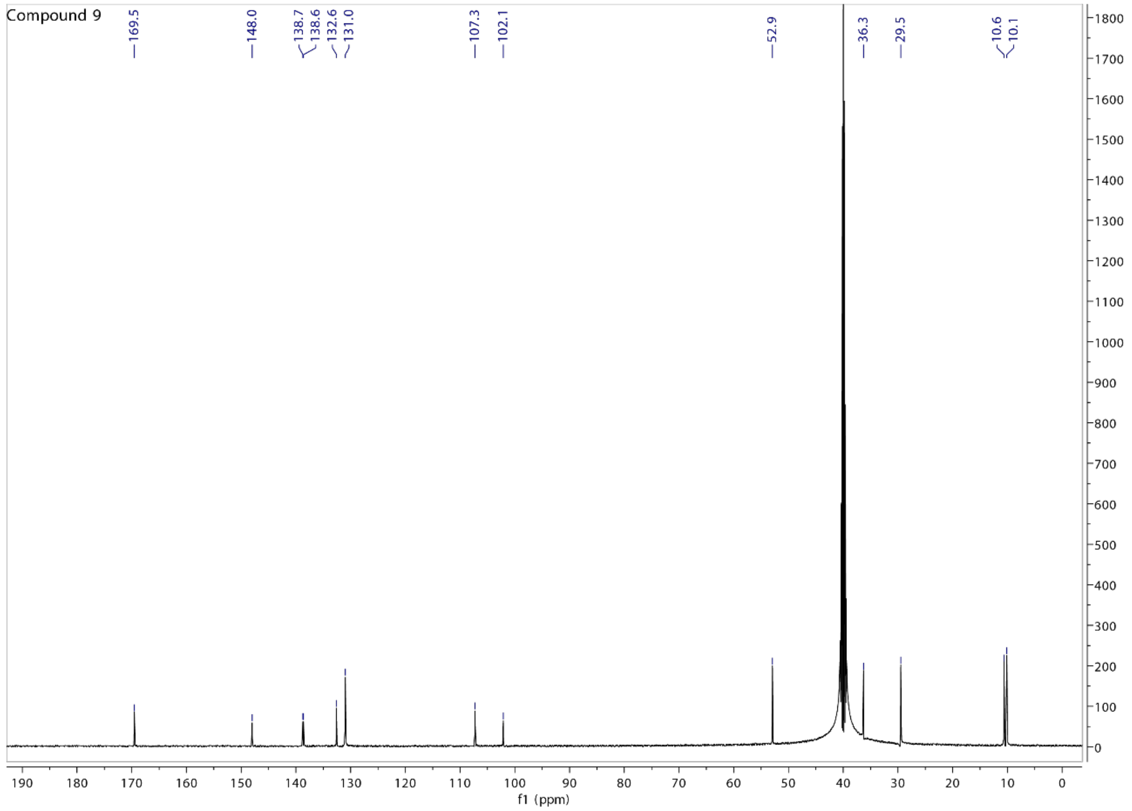
Compound 9
169.5
148.0
138.7
138.6
132.6
131.0
107.3
102.1
52.9
36.3
29.5
10.6
10.1
f1 (ppm)

Compound 10

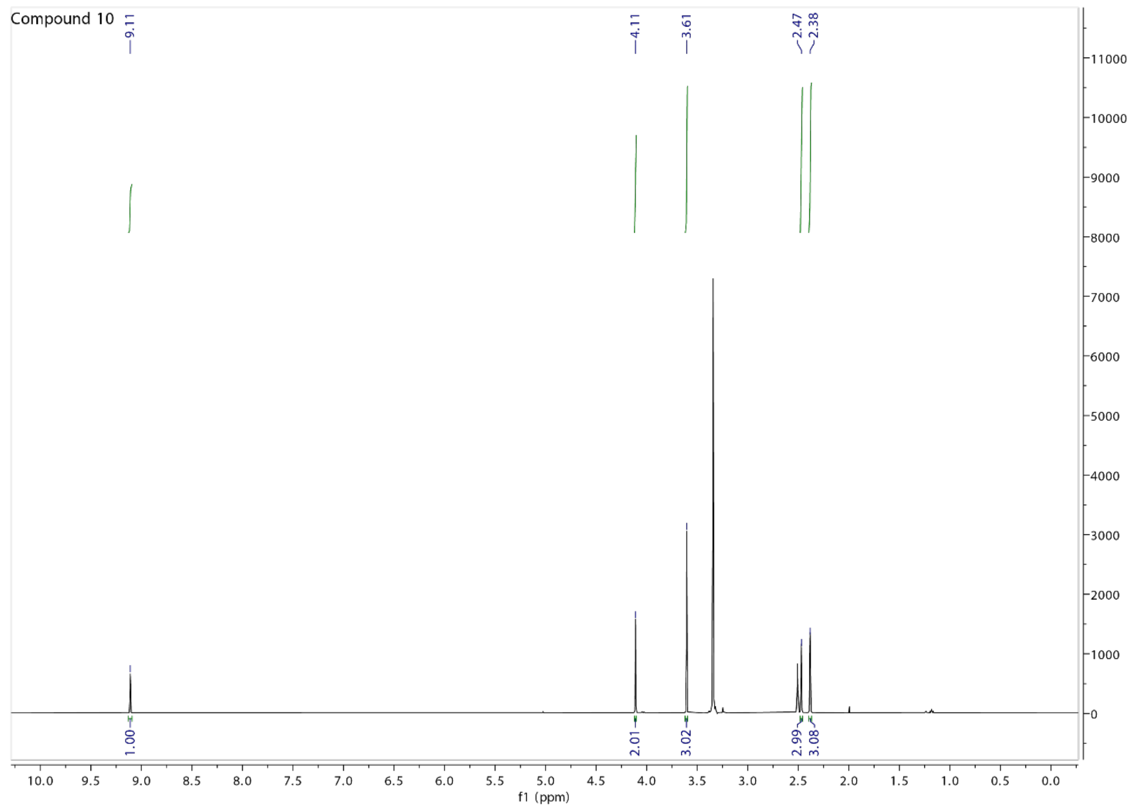

Compound 10
9.11
4.11
3.61
2.47
2.38
1.00
2.01
3.02
2.99
3.08
f1 (ppm)

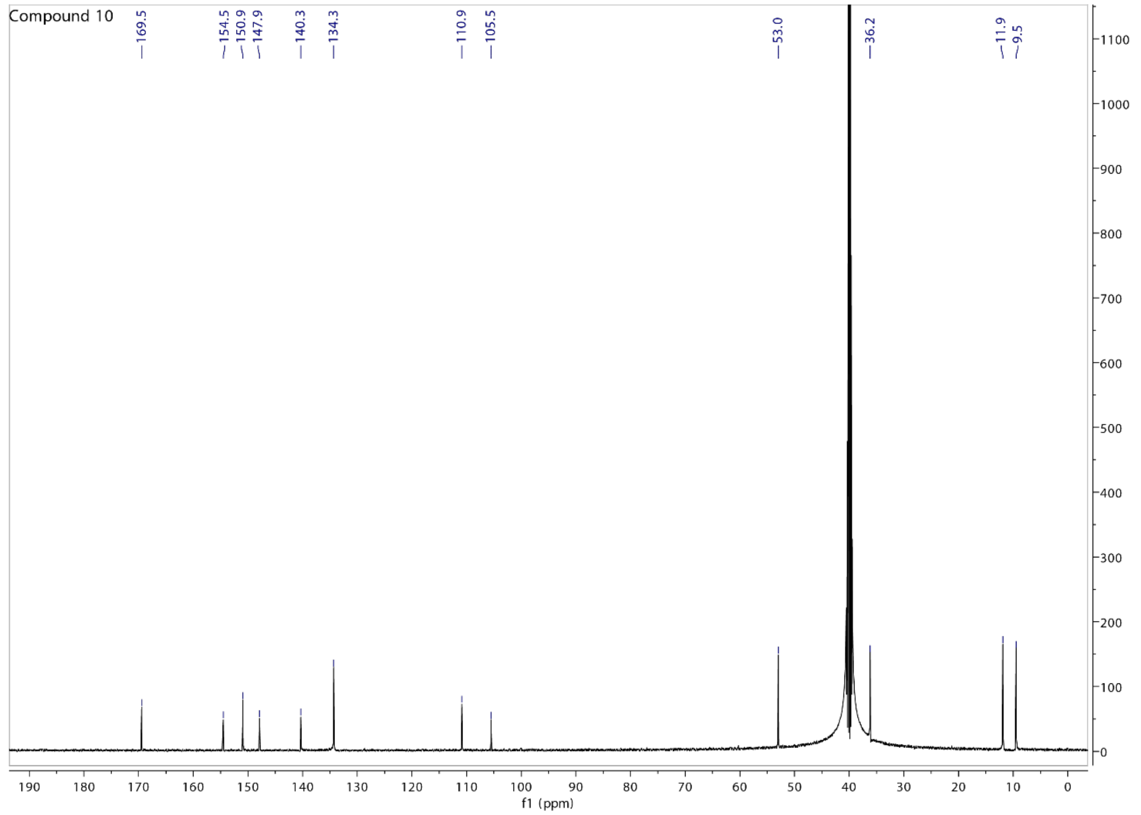

Compound 10
169.5
154.5
150.9
147.9
140.3
134.3
110.9
105.5
53.0
36.2
11.9
9.5
f1 (ppm)

Compound 11

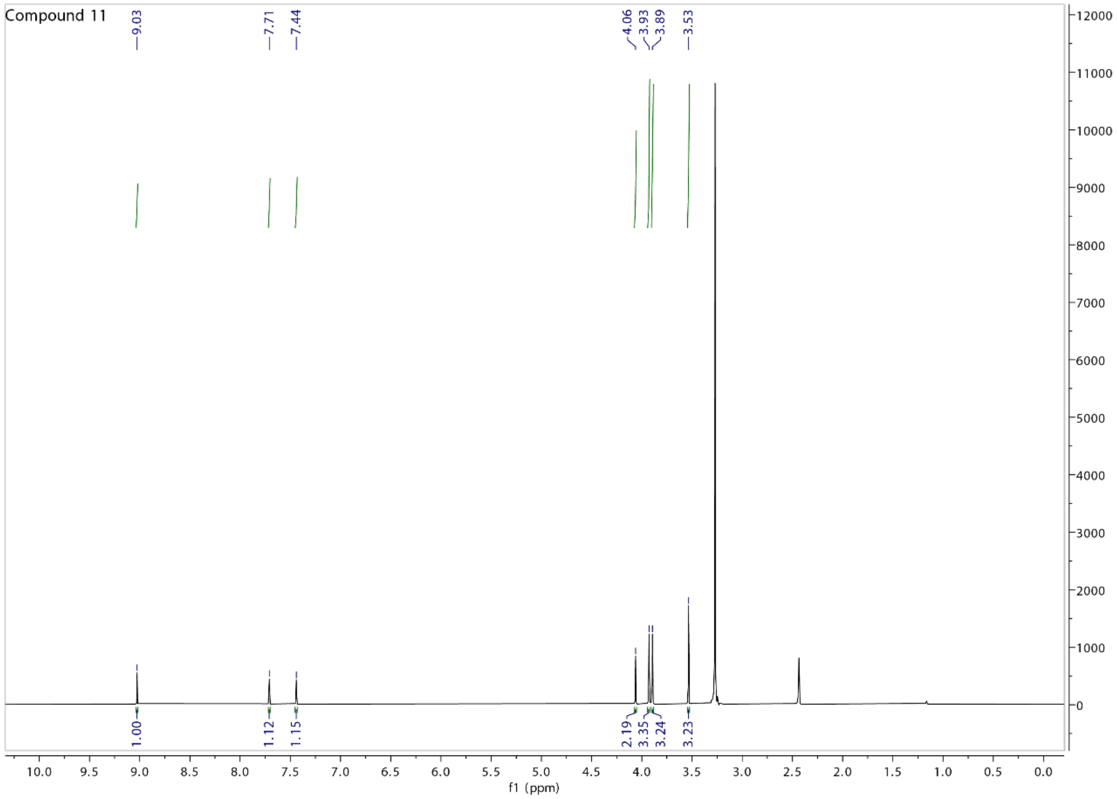
Compound 11
9.03
7.71
7.44
4.06
3.93
3.89
3.53
1.00
1.12
1.15
2.19
3.35
3.24
3.23
f1 (ppm)

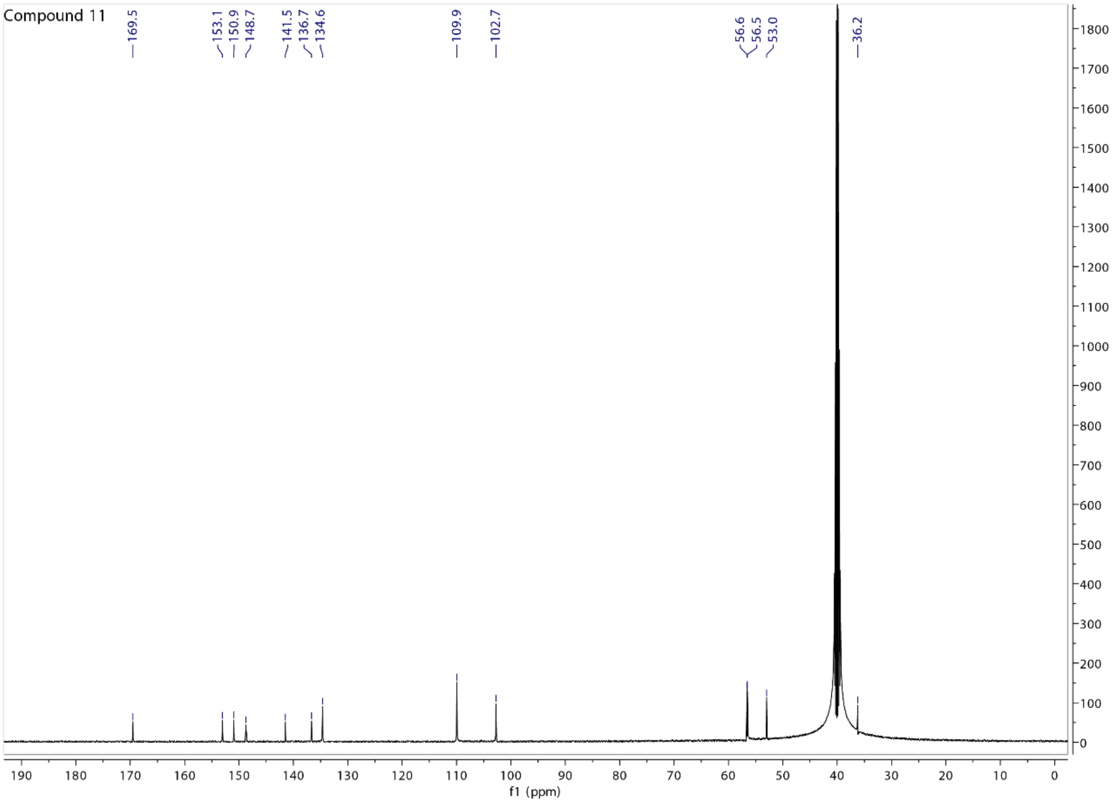
Compound 11
169.5
153.1
150.9
148.7
141.5
136.7
134.6
109.9
102.7
56.6
56.5
53.0
36.2
f1 (ppm)

### Compound 15

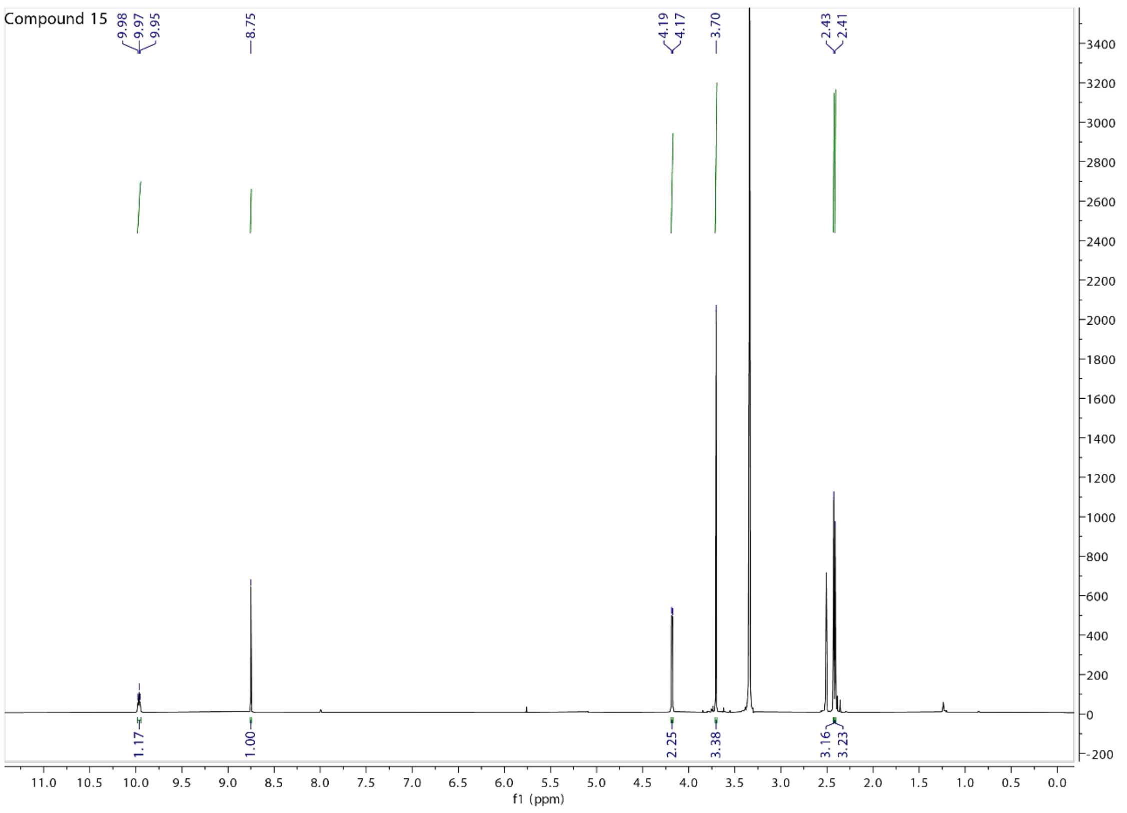
Compound 15
9.98
9.97
9.95
8.75
4.19
4.17
3.70
2.43
2.41
1.17
1.00
2.25
3.38
3.16
3.23
f1 (ppm)

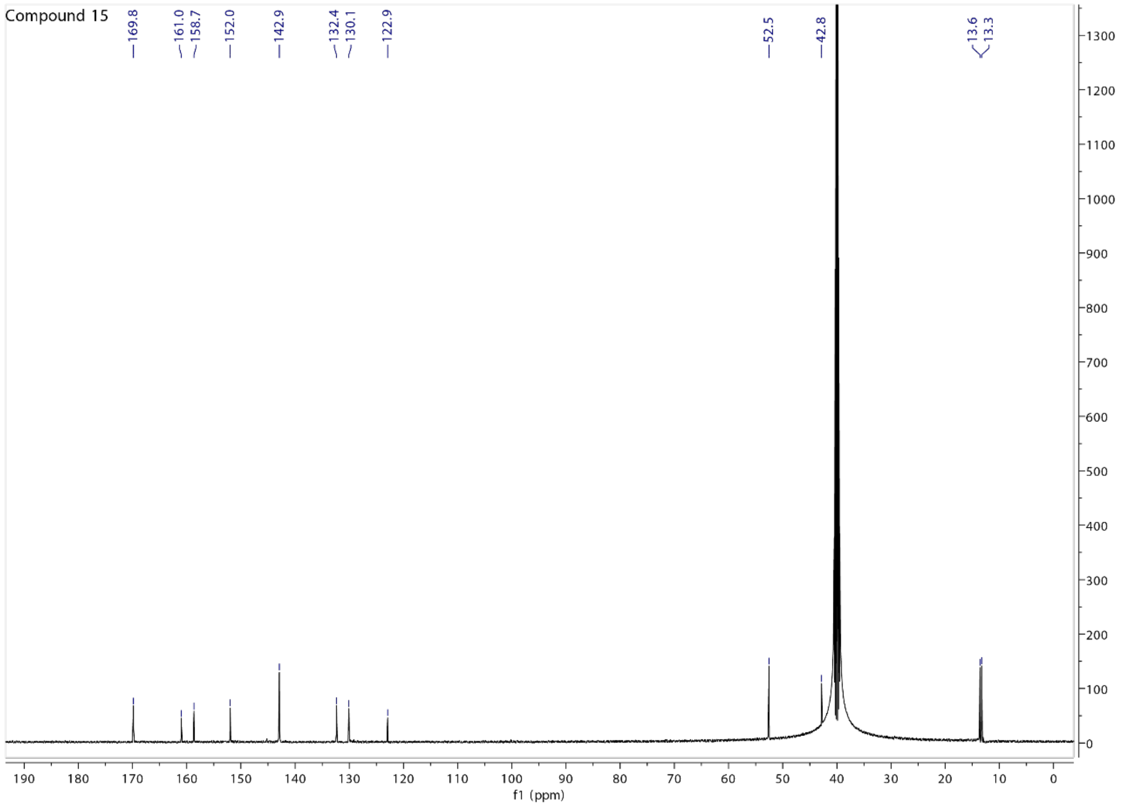
Compound 15
169.8
161.0
158.7
152.0
142.9
132.4
130.1
122.9
52.5
42.8
13.6
13.3
f1 (ppm)

Compound 16

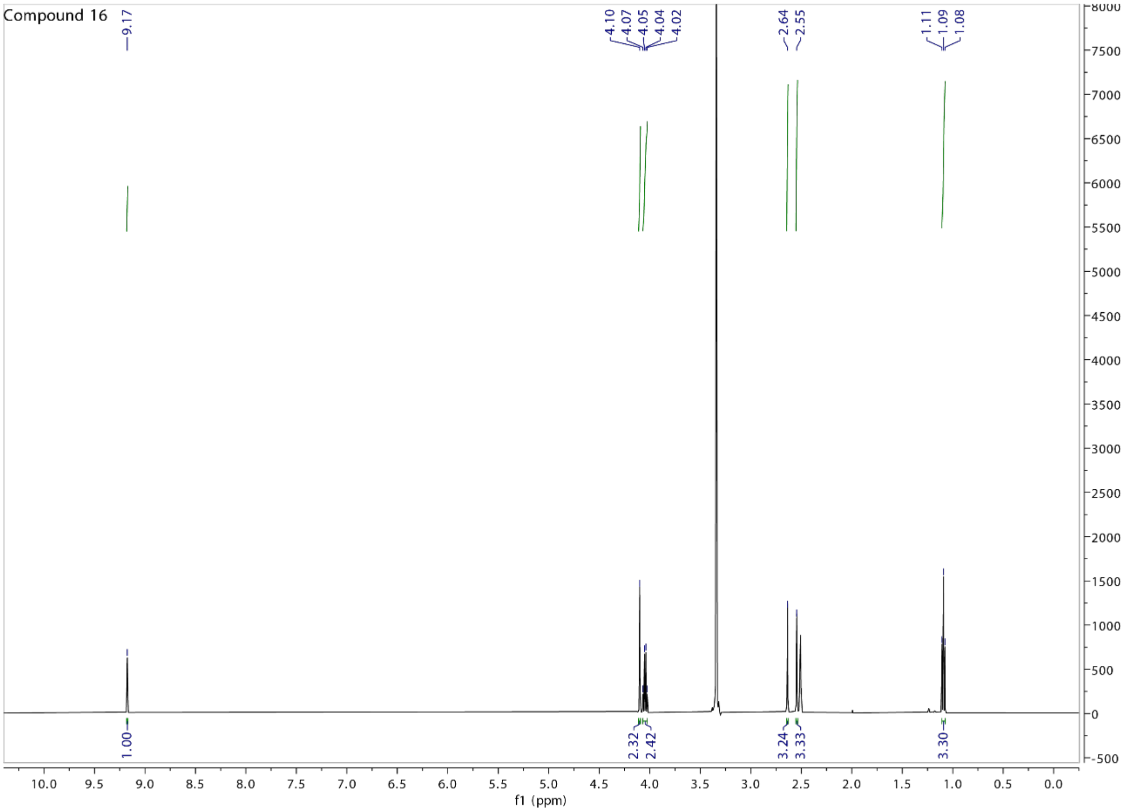

Compound 16
9.17
4.10
4.07
4.05
4.04
4.02
2.64
2.55
1.11
1.09
1.08
1.00
2.32
2.42
3.24
3.33
3.30
f1 (ppm)

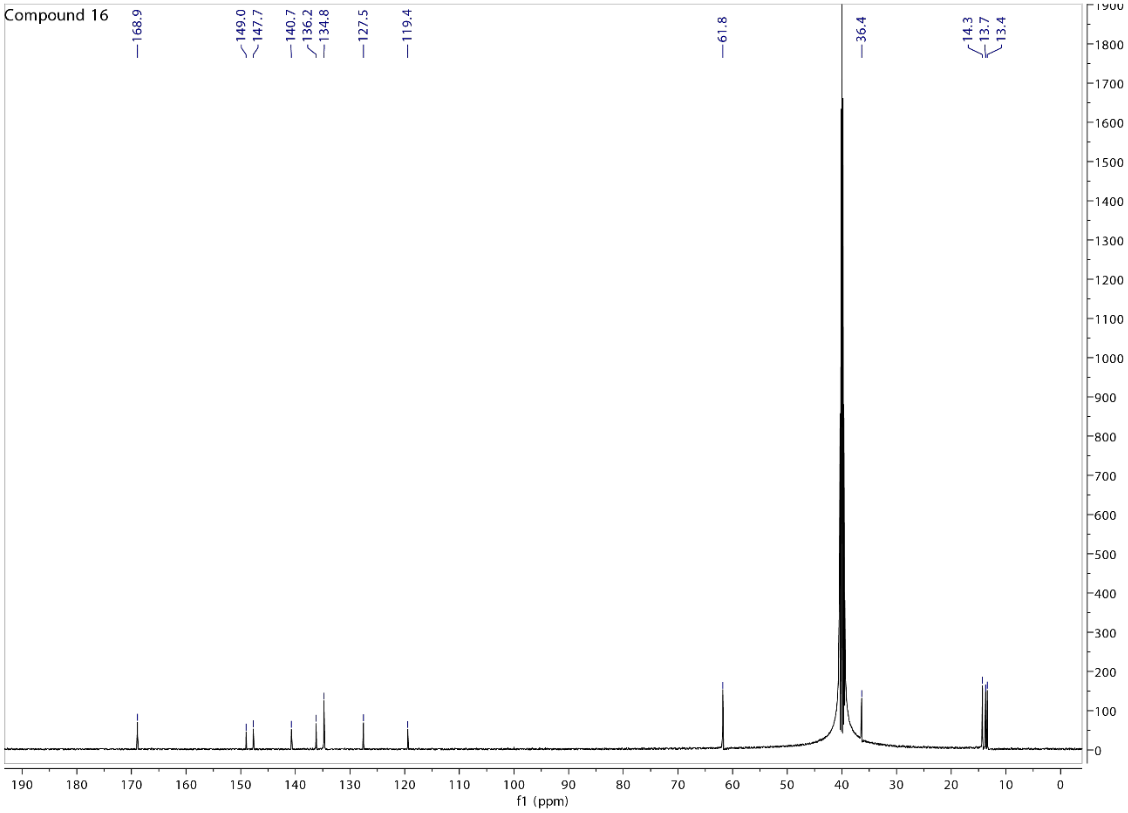

Compound 16
168.9
149.0
147.7
140.7
136.2
134.8
127.5
119.4
61.8
36.4
14.3
13.7
13.4
f1 (ppm)

Compound 19

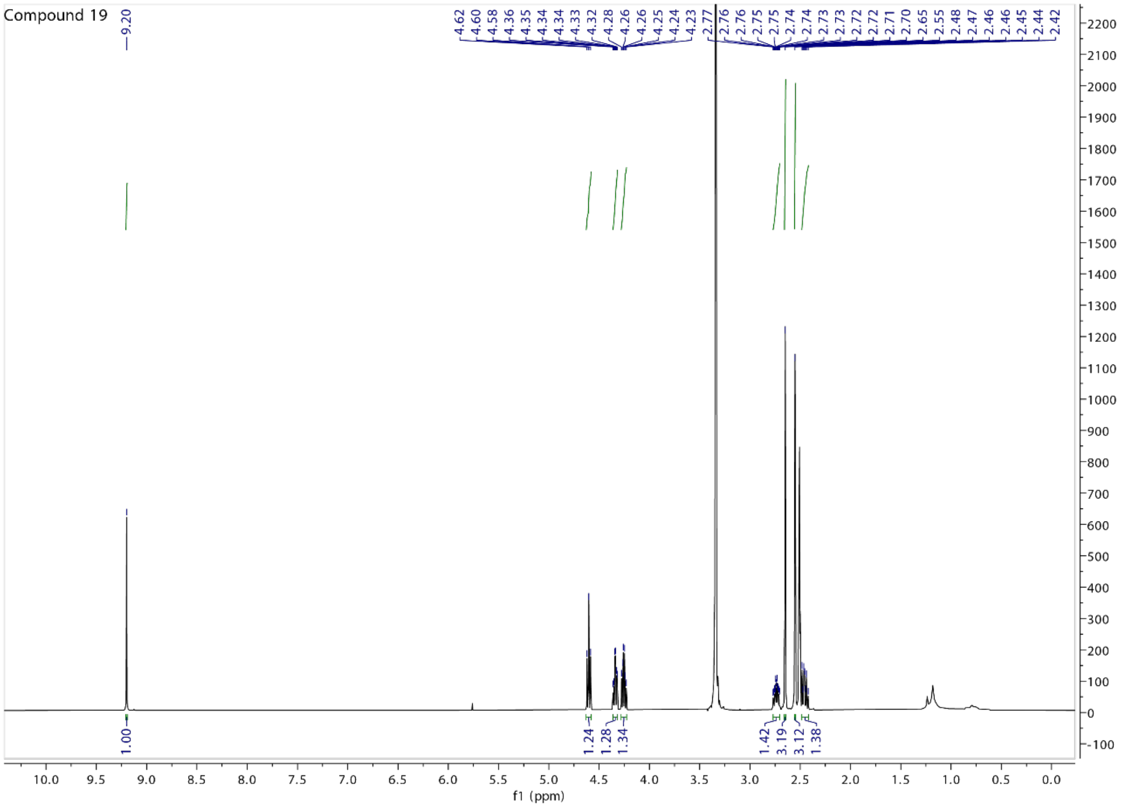
Compound 19
9.20
4.62 4.60 4.58 4.36 4.35 4.34 4.34 4.33 4.32 4.28 4.26 4.26 4.25 4.24 4.23
2.77 2.76 2.76 2.75 2.75 2.74 2.74 2.74 2.73 2.73 2.72 2.72 2.72 2.71 2.70 2.65 2.55 2.48 2.47 2.46 2.46 2.45 2.44 2.42
1.00
1.24 1.28 1.34
1.42 3.19 3.12 1.38
f1 (ppm)

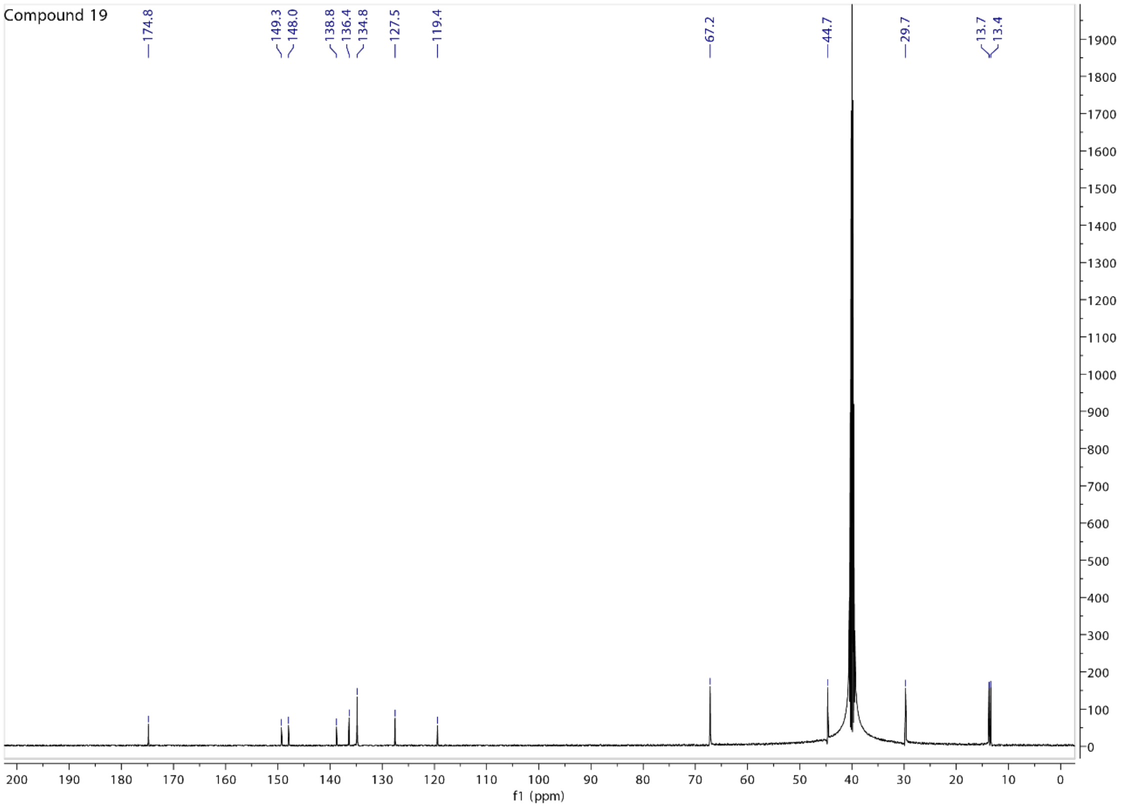
Compound 19
174.8
149.3 148.0
138.8 136.4 134.8
127.5
119.4
67.2
44.7
29.7
13.7 13.4
f1 (ppm)

Compound 21

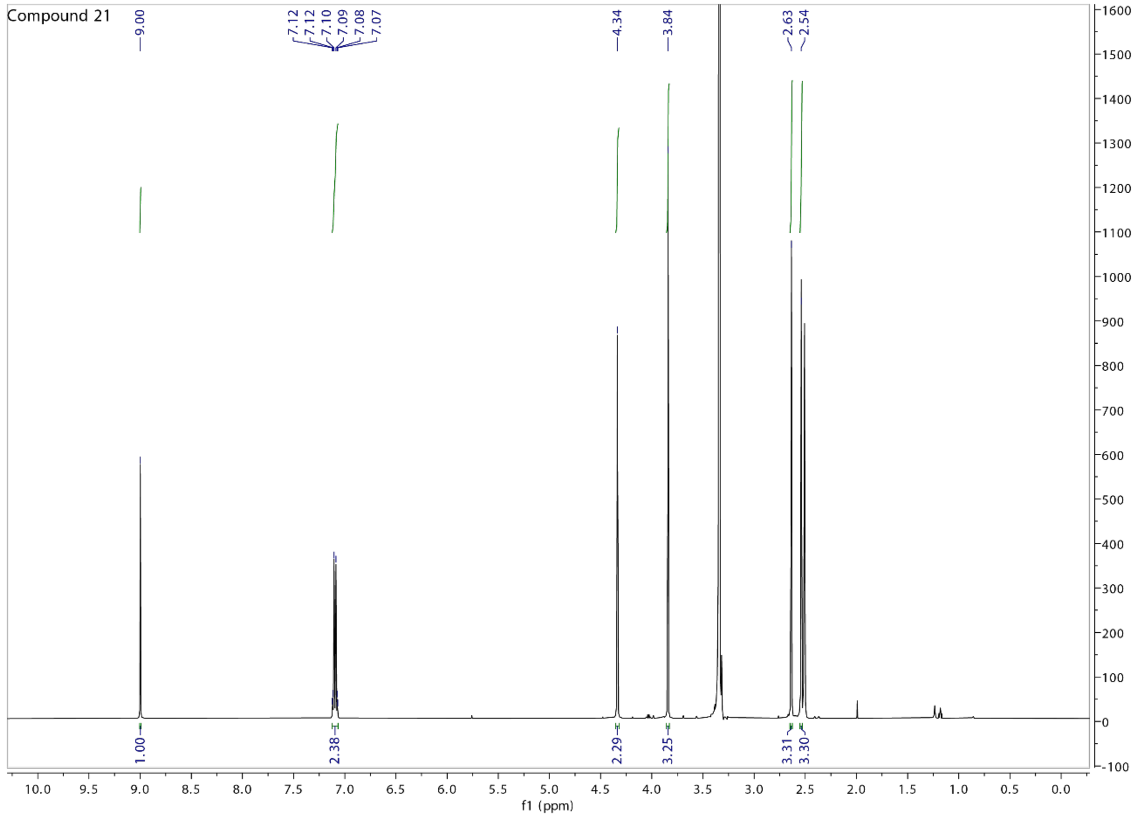
Compound 21
9.00
7.12 7.12 7.10 7.09 7.08 7.07
4.34
3.84
2.63 2.54
1.00
2.38
2.29
3.25
3.31 3.30
f1 (ppm)

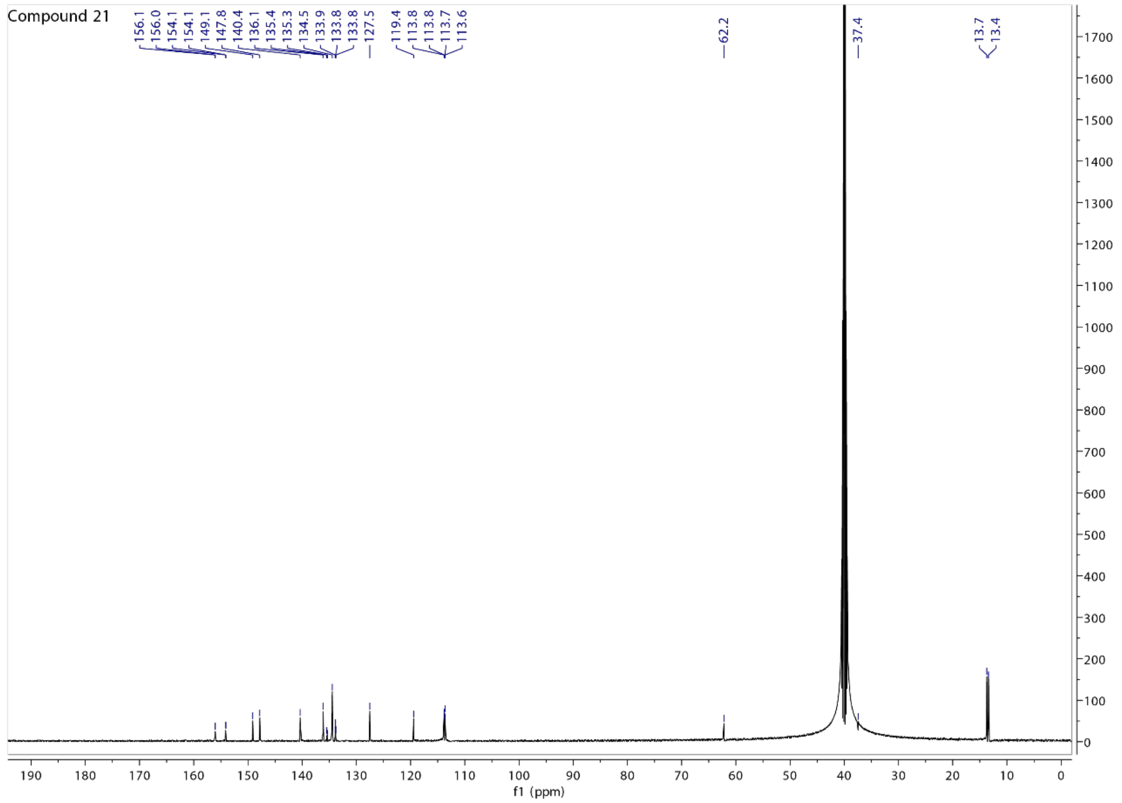
Compound 21
156.1 156.0 154.1 154.1 149.1 147.8 140.4 136.1 135.4 135.3 134.5 133.9 133.8 133.8 127.5 119.4 113.8 113.8 113.7 113.6
62.2
37.4
13.7 13.4
f1 (ppm)

Compound 22

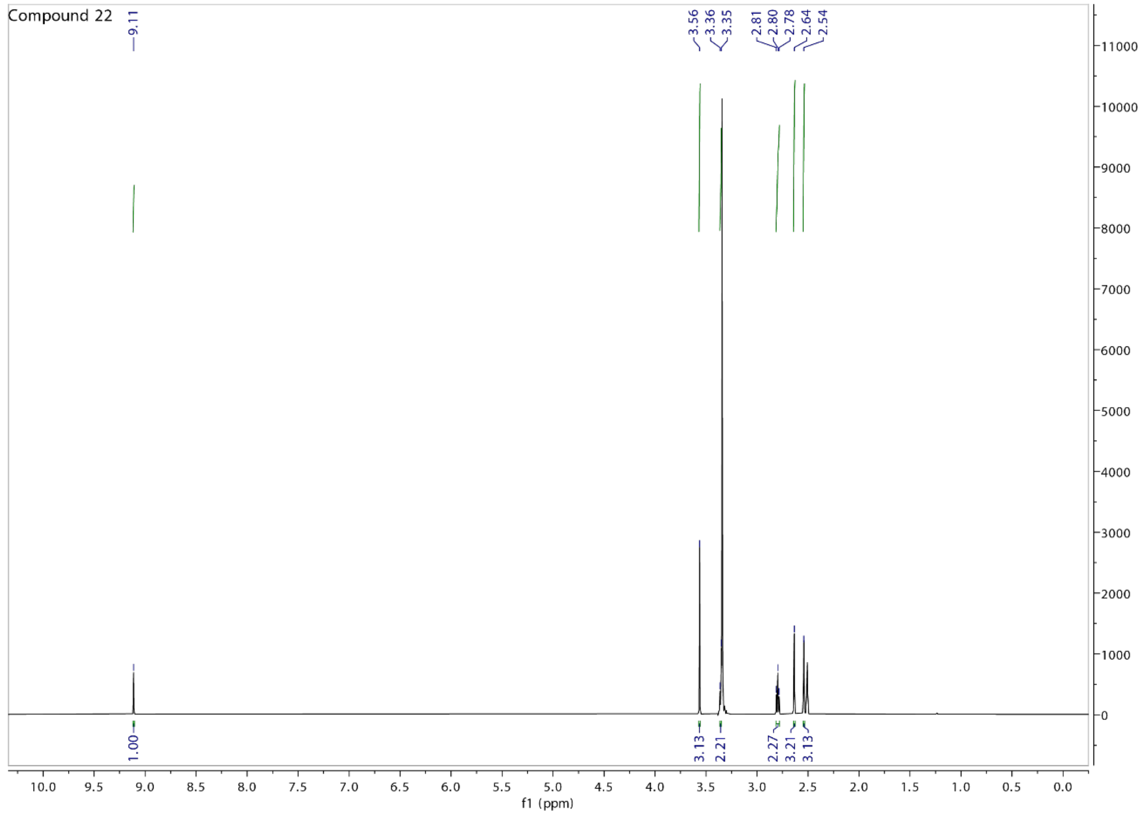
Compound 22
9.11
3.56
3.36
3.35
2.81
2.80
2.78
2.64
2.54
1.00
3.13
2.21
2.27
3.21
3.13
f1 (ppm)

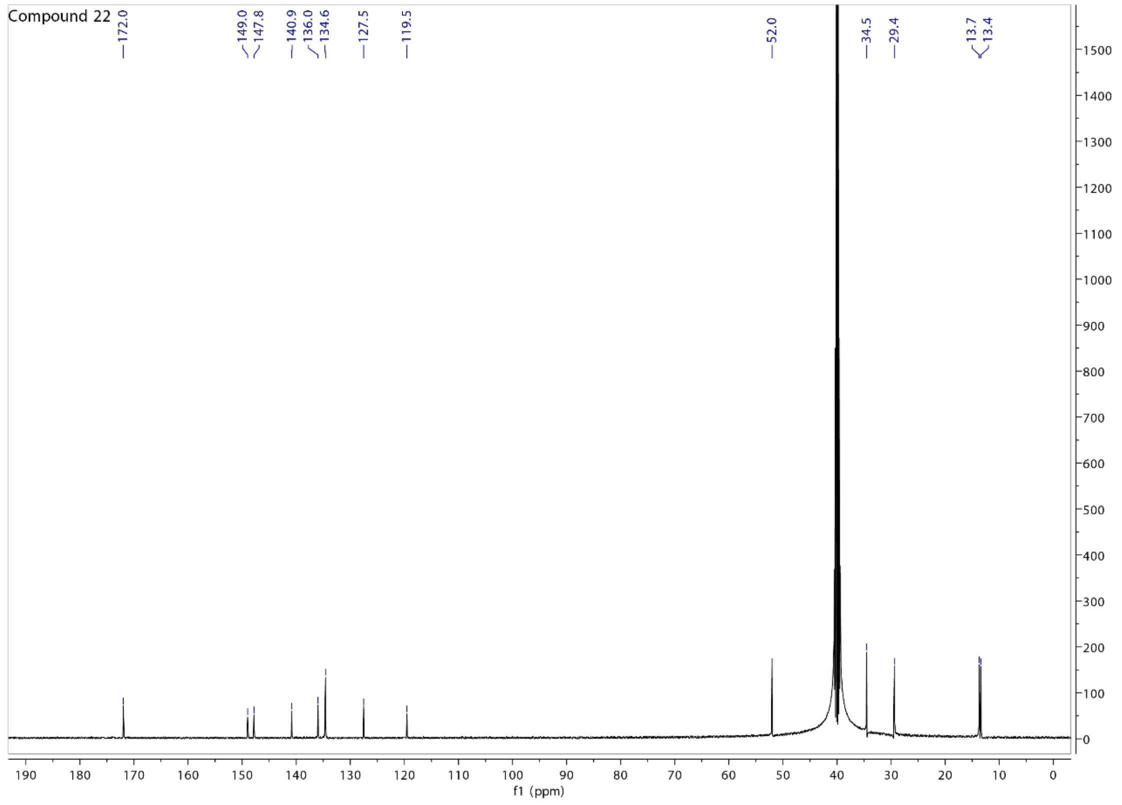
Compound 22
172.0
149.0
147.8
140.9
136.0
134.6
127.5
119.5
52.0
34.5
29.4
13.7
13.4
f1 (ppm)

Compound 23

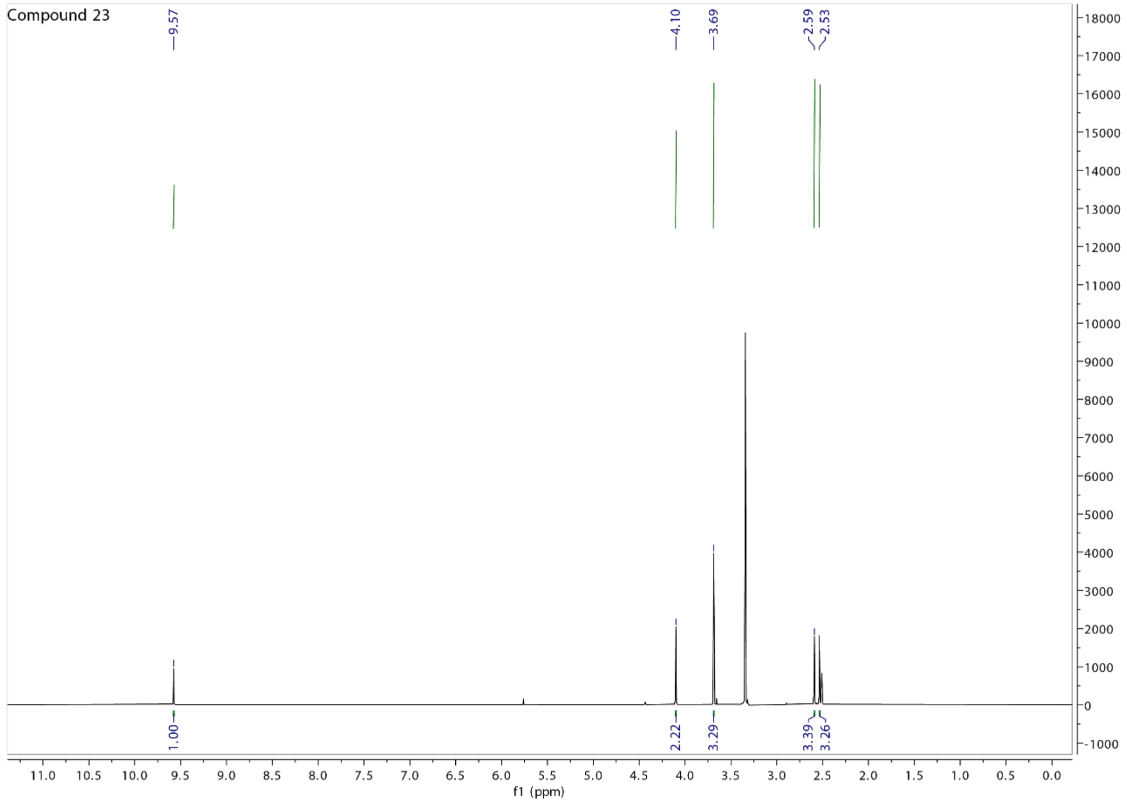
Compound 23
9.57
4.10
3.69
2.59
2.53
1.00
2.22
3.29
3.39
3.26
f1 (ppm)

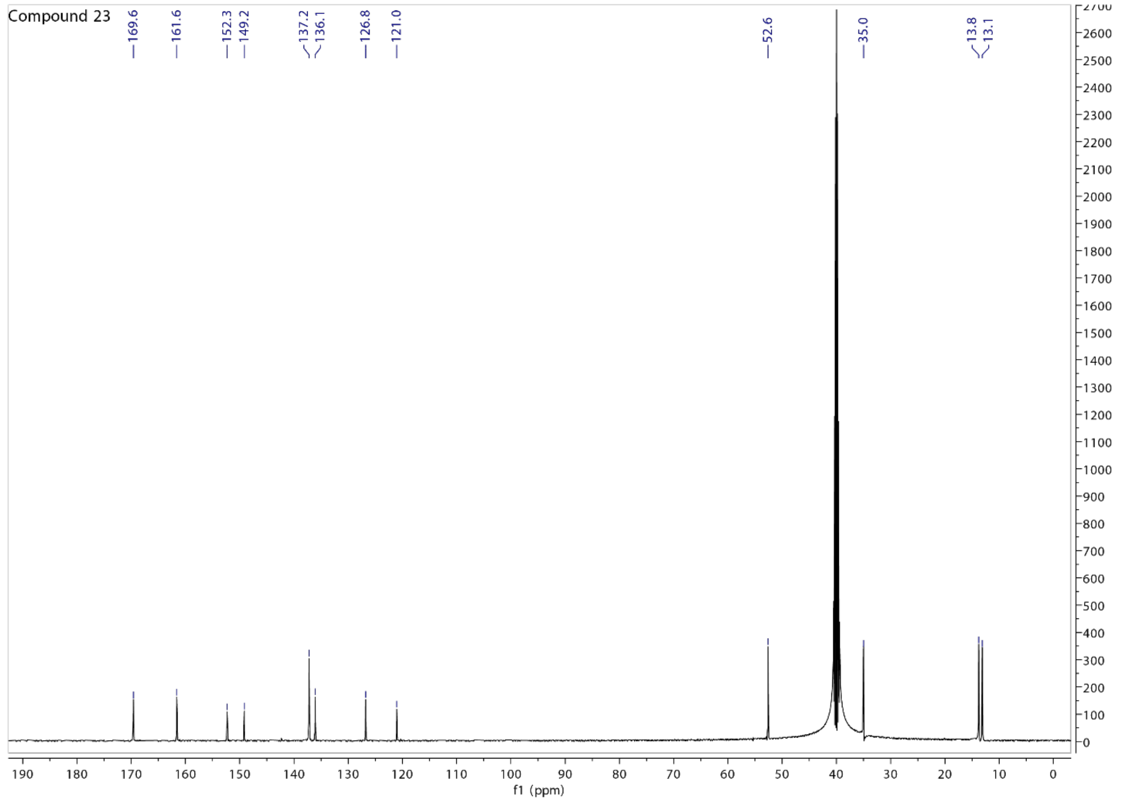
Compound 23
169.6
161.6
152.3
149.2
137.2
136.1
126.8
121.0
52.6
35.0
13.8
13.1
f1 (ppm)

Compound 24

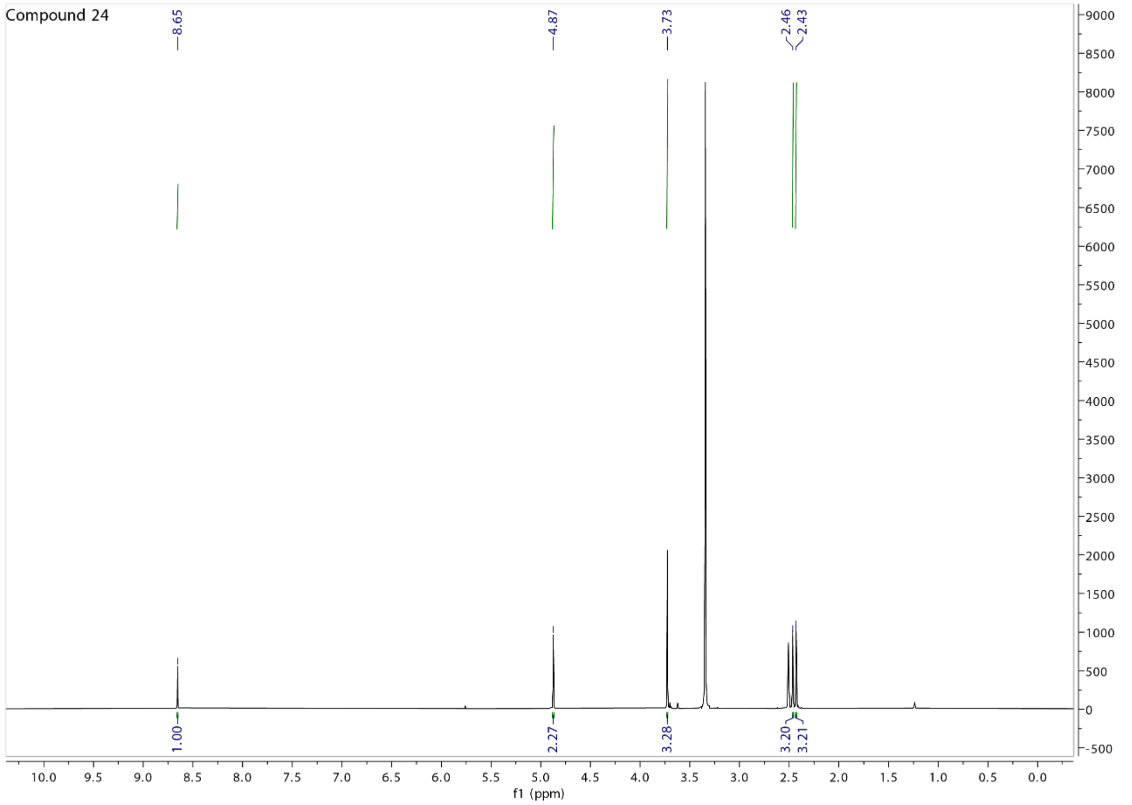
Compound 24
8.65
4.87
3.73
2.46
2.43
1.00
2.27
3.28
3.20
3.21
f1 (ppm)

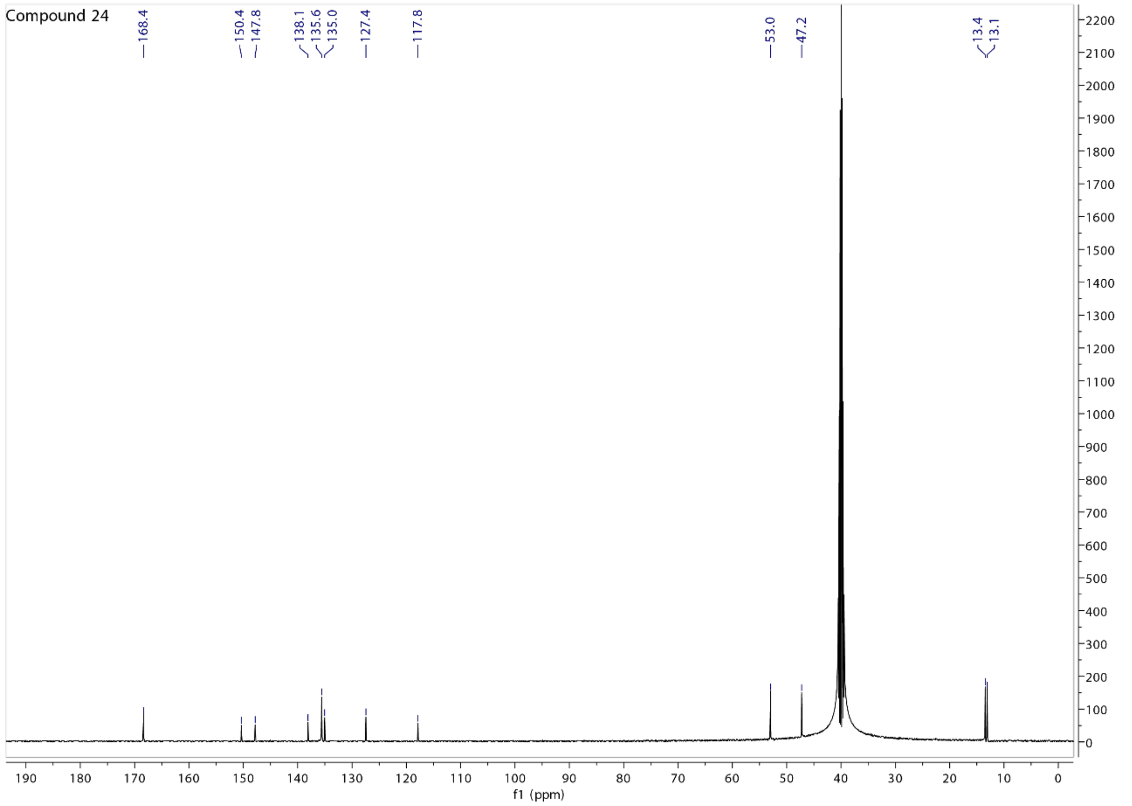
Compound 24
168.4
150.4
147.8
138.1
135.6
135.0
127.4
117.8
53.0
47.2
13.4
13.1
f1 (ppm)

Compound 25

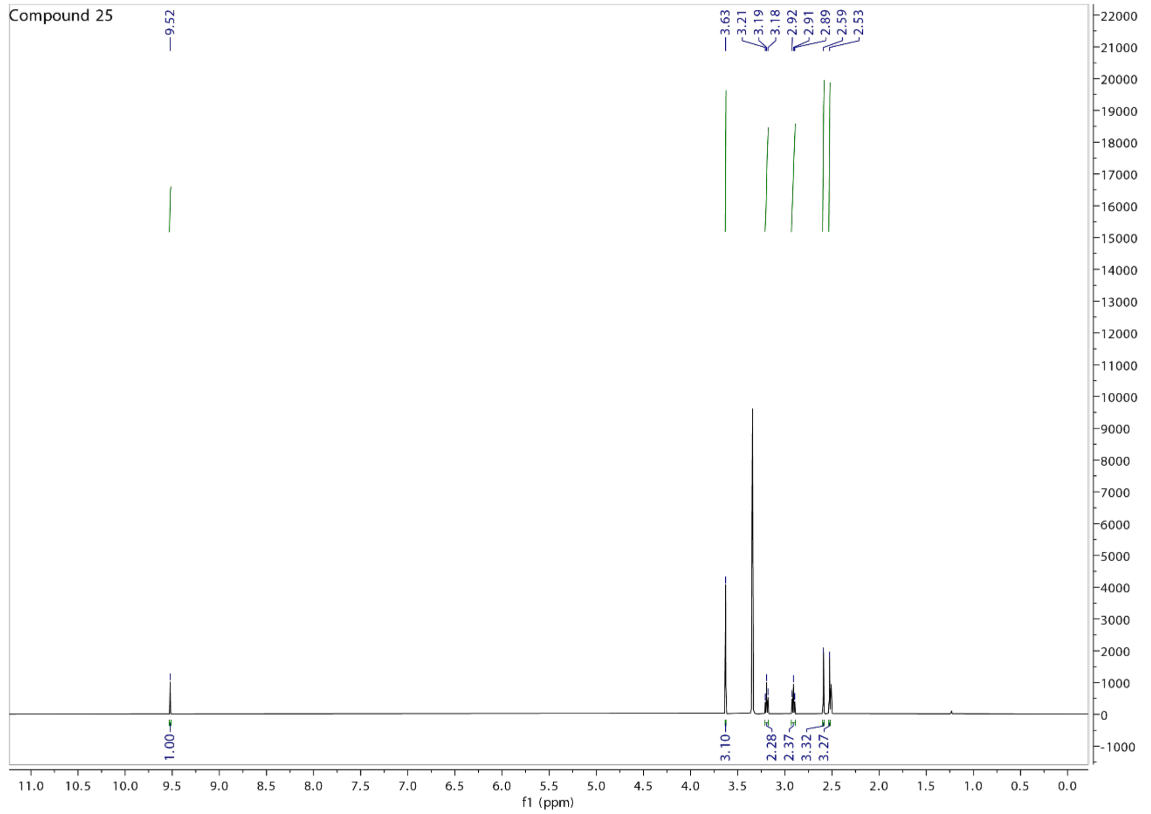
Compound 25
9.52
3.63
3.21
3.19
3.18
2.92
2.91
2.89
2.59
2.53
1.00
3.10
2.28
2.37
3.32
3.27
f1 (ppm)

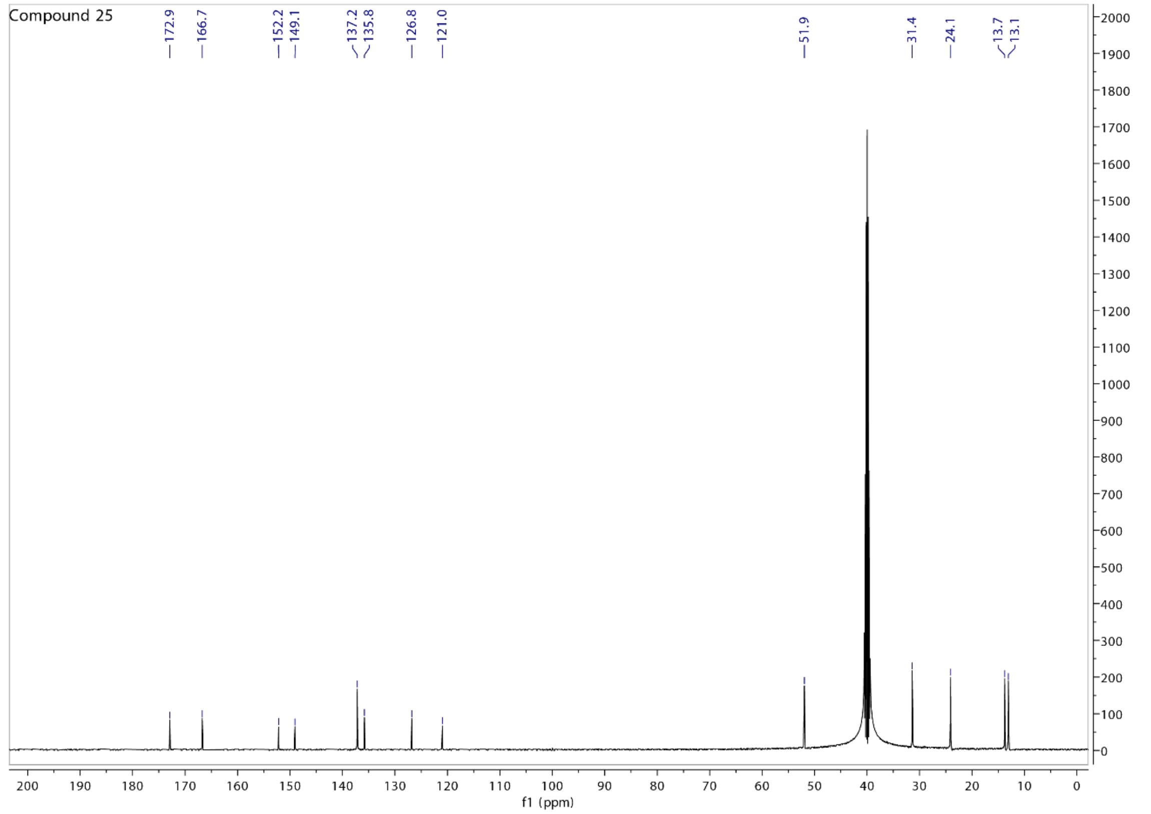
Compound 25
172.9
166.7
152.2
149.1
137.2
135.8
126.8
121.0
51.9
31.4
24.1
13.7
13.1
f1 (ppm)

Compound 26

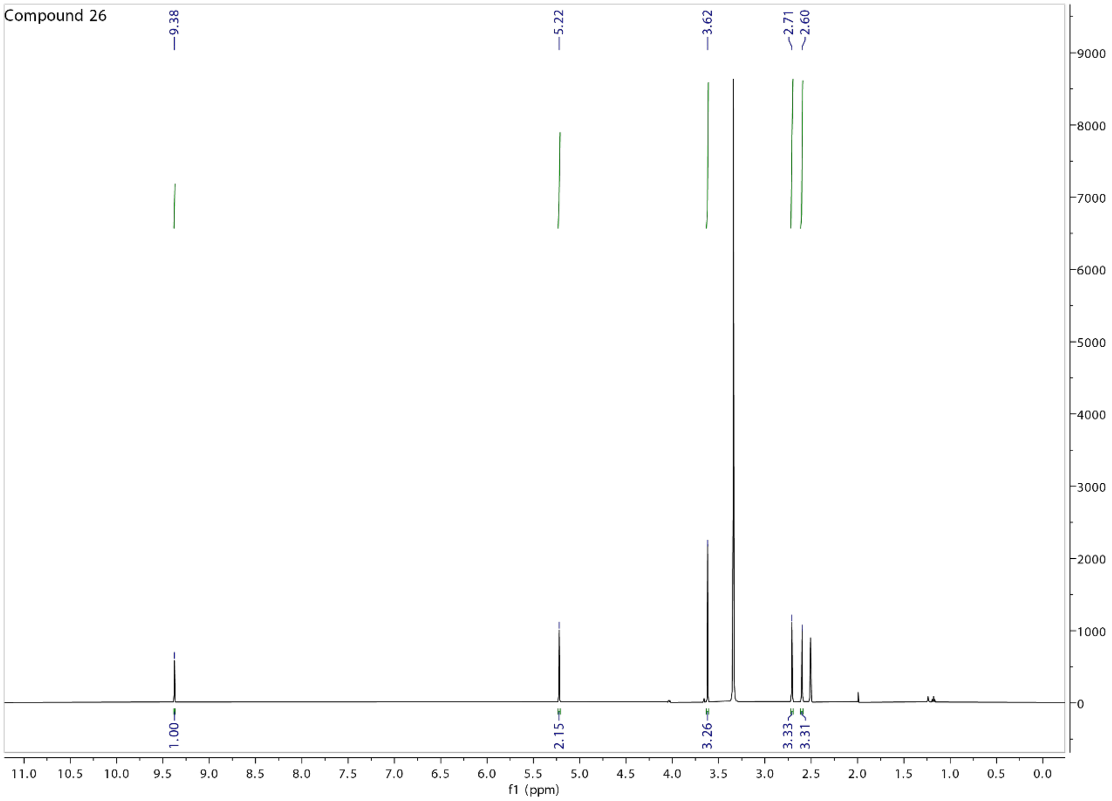
Compound 26
9.38
5.22
3.62
2.71
2.60
1.00
2.15
3.26
3.33
3.31
f1 (ppm)

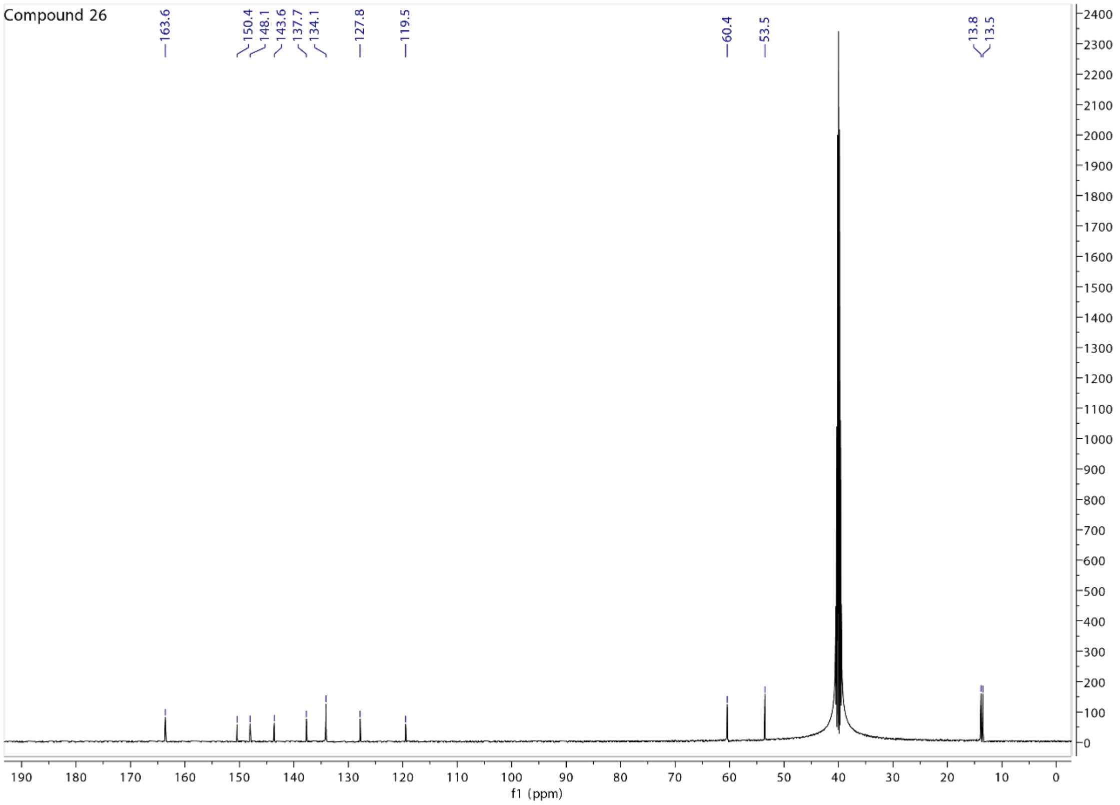
Compound 26
163.6
150.4
148.1
143.6
137.7
134.1
127.8
119.5
60.4
53.5
13.8
13.5
f1 (ppm)

Purity analysis of active compounds **BDW568**, **1**, **3**, **6**, **8**, **16** and **17**

Compound BDW568

Empower 3 — Default Individual Report

SAMPLE INFORMATION

| | | | |
|---|---|---|---|
| Sample Name: | zt-bdw568 | Acquired By: | System |
| Sample Type: | Unknown | Sample Set Name: | ztsm |
| Vial: | 1:E,7 | Acq. Method Set: | L4ABT1 |
| Injection #: | 1 | Processing Method: | bdw568 |
| Injection Volume: | 3.00 ul | Channel Name: | PDA Ch1 254nm@4.8nm |
| Run Time: | 5.0 Minutes | Proc. Chnl. Descr.: | PDA Ch1 254nm@4.8nm |
| Date Acquired: | 6/21/2023 9:03:26 PM CDT | | |
| Date Processed: | 6/24/2023 6:08:31 PM CDT | | |

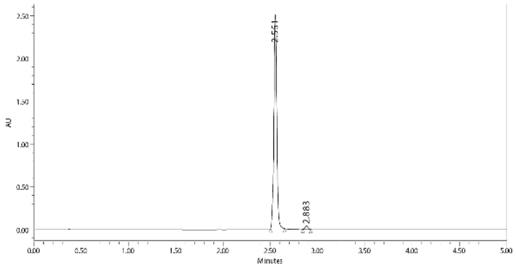

| | RT | Area | % Area | Height |
|---|---|---|---|---|
| 1 | 2.551 | 5332833 | 98.36 | 2502237 |
| 2 | 2.883 | 88891 | 1.64 | 40562 |

Reported by User: System
Report Method: Default Individual Report
Report Method ID 1003
Page: 1 of 1

Project Name: UPLC
Date Printed:
6/24/2023
6:08:48 PM US/Central

Compound 1

Empower 3 — Default Individual Report

SAMPLE INFORMATION

| | | | |
|---|---|---|---|
| Sample Name: | zt-bdw-jmc-1 | Acquired By: | System |
| Sample Type: | Unknown | Sample Set Name: | ztsm |
| Vial: | 1:E,1 | Acq. Method Set: | L4ABT1 |
| Injection #: | 1 | Processing Method: | bdwjmc1 |
| Injection Volume: | 5.00 ul | Channel Name: | PDA Ch1 254nm@4.8nm |
| Run Time: | 5.0 Minutes | Proc. Chnl. Descr.: | PDA Ch1 254nm@4.8nm |
| Date Acquired: | 6/21/2023 8:23:14 PM CDT | | |
| Date Processed: | 6/24/2023 4:38:59 PM CDT | | |

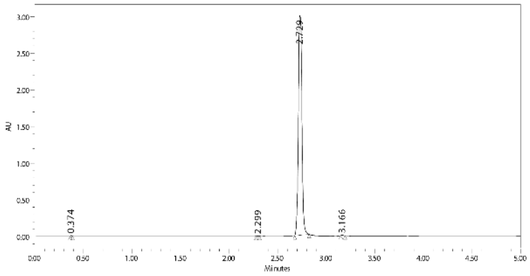

| | RT | Area | % Area | Height |
|---|---|---|---|---|
| 1 | 0.374 | 10846 | 0.14 | 13884 |
| 2 | 2.299 | 3810 | 0.05 | 2983 |
| 3 | 2.729 | 7649022 | 99.70 | 2998439 |
| 4 | 3.166 | 8088 | 0.11 | 5345 |

Reported by User: System
Report Method: Default Individual Report
Report Method ID 1003
Page: 1 of 1

Project Name: UPLC
Date Printed:
6/24/2023
4:39:55 PM US/Central

#### Compound 3

Empower 3

Default Individual Report

SAMPLE INFORMATION

| | | | |
|---|---|---|---|
| Sample Name: | zt-bdw-jmc-3 | Acquired By: | System |
| Sample Type: | Unknown | Sample Set Name: | ztsm |
| Vial: | 1:E,3 | Acq. Method Set: | L4ABT1 |
| Injection #: | 1 | Processing Method: | bdwjmc3 |
| Injection Volume: | 3.00 ul | Channel Name: | PDA Ch1 254nm@4.8nm |
| Run Time: | 5.0 Minutes | Proc. Chnl. Descr.: | PDA Ch1 254nm@4.8nm |
| Date Acquired: | 6/21/2023 8:40:56 PM CDT | | |
| Date Processed: | 6/24/2023 4:42:31 PM CDT | | |

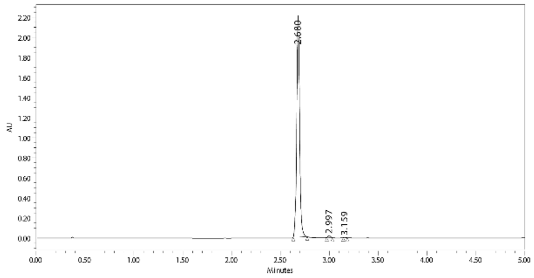

| | RT | Area | % Area | Height |
|---|---|---|---|---|
| 1 | 2.680 | 4620866 | 99.13 | 2207163 |
| 2 | 2.997 | 34228 | 0.73 | 20367 |
| 3 | 3.159 | 6413 | 0.14 | 4594 |

Reported by User: System
Report Method: Default Individual Report
Report Method ID 1003
Page: 1 of 1

Project Name: UPLC
Date Printed:
6/24/2023
4:42:53 PM US/Central

#### Compound 6

Empower 3

Default Individual Report

SAMPLE INFORMATION

| | | | |
|---|---|---|---|
| Sample Name: | zt-bdw-jmc-6 | Acquired By: | System |
| Sample Type: | Unknown | Sample Set Name: | ztsm |
| Vial: | 1:E,6 | Acq. Method Set: | L4ABT1 |
| Injection #: | 1 | Processing Method: | bdwjmc6 |
| Injection Volume: | 3.00 ul | Channel Name: | PDA Ch1 254nm@4.8nm |
| Run Time: | 5.0 Minutes | Proc. Chnl. Descr.: | PDA Ch1 254nm@4.8nm |
| Date Acquired: | 6/21/2023 8:57:49 PM CDT | | |
| Date Processed: | 6/24/2023 4:44:13 PM CDT | | |

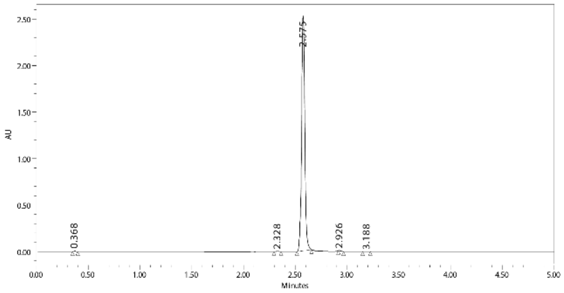

| | RT | Area | % Area | Height |
|---|---|---|---|---|
| 1 | 0.368 | 7467 | 0.14 | 9720 |
| 2 | 2.328 | 7697 | 0.14 | 4407 |
| 3 | 2.575 | 5396829 | 99.62 | 2524547 |
| 4 | 2.926 | 4245 | 0.08 | 3085 |
| 5 | 3.188 | 1348 | 0.02 | -634 |

Reported by User: System
Report Method: Default Individual Report
Report Method ID 1003
Page: 1 of 1

Project Name: UPLC
Date Printed:
6/24/2023
4:44:31 PM US/Central

### Compound 8

Empower 3

#### Default Individual Report

##### SAMPLE INFORMATION

| | | | |
|---|---|---|---|
| Sample Name: | zt-bdw-jmc-8 | Acquired By: | System |
| Sample Type: | Unknown | Sample Set Name: | ztsm |
| Vial: | 1:E,8 | Acq. Method Set: | L4ABT1 |
| Injection #: | 1 | Processing Method: | bdwjmc8 |
| Injection Volume: | 3.00 ul | Channel Name: | PDA Ch1 254nm@4.8nm |
| Run Time: | 5.0 Minutes | Proc. Chnl. Descr.: | PDA Ch1 254nm@4.8nm |
| Date Acquired: | 6/21/2023 9:09:07 PM CDT | | |
| Date Processed: | 6/24/2023 4:45:24 PM CDT | | |

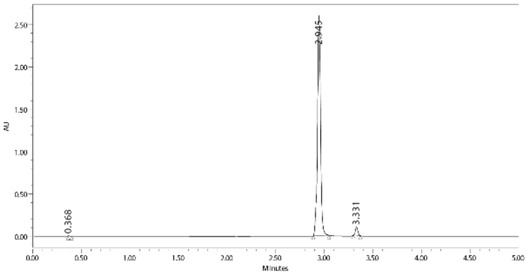

| | RT | Area | % Area | Height |
|---|---|---|---|---|
| 1 | 0.368 | 6010 | 0.10 | 8504 |
| 2 | 2.945 | 5669591 | 96.95 | 2600174 |
| 3 | 3.331 | 172548 | 2.95 | 97772 |

Reported by User: System
Report Method: Default Individual Report
Report Method ID 1003
Page: 1 of 1

Project Name: UPLC
Date Printed:
6/24/2023
4:45:40 PM US/Central

### Compound 16

Empower 3

#### Default Individual Report

##### SAMPLE INFORMATION

| | | | |
|---|---|---|---|
| Sample Name: | zt-bdw-jmc-16 | Acquired By: | System |
| Sample Type: | Unknown | Sample Set Name: | ztsm |
| Vial: | 1:F,2 | Acq. Method Set: | L4ABT1 |
| Injection #: | 1 | Processing Method: | bdwjmc16 |
| Injection Volume: | 5.00 ul | Channel Name: | PDA Ch1 254nm@4.8nm |
| Run Time: | 5.0 Minutes | Proc. Chnl. Descr.: | PDA Ch1 254nm@4.8nm |
| Date Acquired: | 6/21/2023 3:45:10 PM CDT | | |
| Date Processed: | 6/24/2023 6:11:05 PM CDT | | |

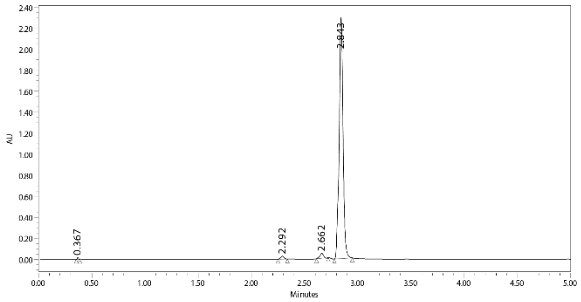

| | RT | Area | % Area | Height |
|---|---|---|---|---|
| 1 | 0.367 | 13319 | 0.22 | 17455 |
| 2 | 2.292 | 59641 | 0.97 | 27426 |
| 3 | 2.662 | 113402 | 1.85 | 46986 |
| 4 | 2.843 | 5944465 | 96.96 | 2297714 |

Reported by User: System
Report Method: Default Individual Report
Report Method ID 1003
Page: 1 of 1

Project Name: UPLC
Date Printed:
6/24/2023
6:11:26 PM US/Central

Compound 17

Default Individual Report

SAMPLE INFORMATION

| | | | |
|---|---|---|---|
| Sample Name: | zt-bdw-jmc17 | Acquired By: | System |
| Sample Type: | Unknown | Sample Set Name: | ztsm |
| Vial: | 1:E,5 | Acq. Method Set: | L4ABT1 |
| Injection #: | 1 | Processing Method: | bdwjmc17 |
| Injection Volume: | 3.00 ul | Channel Name: | PDA Ch1 254nm@4.8nm |
| Run Time: | 5.0 Minutes | Proc. Chnl. Descr.: | PDA Ch1 254nm@4.8nm |
| Date Acquired: | 6/21/2023 8:52:11 PM CDT | | |
| Date Processed: | 6/24/2023 6:09:54 PM CDT | | |

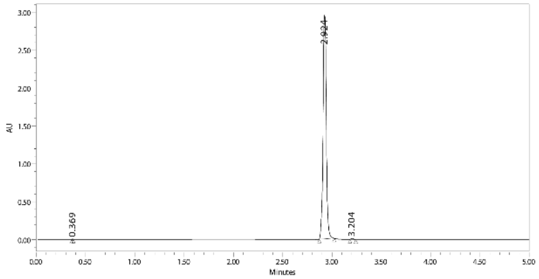

| | RT | Area | % Area | Height |
|---|---|---|---|---|
| 1 | 0.369 | 4827 | 0.07 | 8094 |
| 2 | 2.924 | 6747073 | 99.62 | 2950454 |
| 3 | 3.204 | 20984 | 0.31 | 12909 |
